# Genomic architecture and evolutionary conflict drive allele-specific expression in the social supergene of the red fire ant

**DOI:** 10.1101/2020.02.28.969998

**Authors:** Carlos Martinez-Ruiz, Rodrigo Pracana, Eckart Stolle, Carolina I. Paris, Richard A. Nichols, Yannick Wurm

## Abstract

Supergenes are genomic regions of suppressed recombination that underlie complex polymorphisms. Despite the importance of such regions, our empirical understanding of their early evolution is limited. The young “social” supergene of the fire ant *Solenopsis invicta* provides a powerful system for disentangling the roles of evolutionary conflict and the implications of suppressed recombination.

We used population genomics to identify genetic differences between supergene variants and gene expression analyses across different populations, castes and body parts to characterize allelic expression differences for the hundreds of genes in the supergene.

We find that the expression of most genes is independent of social form or supergene variant, in line with the young age of this system. Many of the genes with allelic expression differences, however, show a pattern consistent with gene degeneration due to suppressed recombination. In contrast, a small portion of the genes has the signature of evolutionary conflict between social forms.

## Introduction

Selection for multiple phenotypic optima within a single species can result in intra-specific evolutionary conflict. Such conflict can lead to selection for reduced recombination between co-adapted alleles encoding the different phenotypes (1). In turn, this selection pressure can favor the formation of supergene regions containing tightly linked alleles of up to hundreds of genes. Such regions enable the maintenance of genetic interactions over evolutionary time (2, 3). We now know that supergenes controlling ecologically important traits are widespread. These include flower heterostyly in *Primula (4)*, mating type in *Mycrobotryum* fungi (5), Batesian mimicry in butterflies (6, 7), mating behavior in white-throated sparrows (8, 9) or male sexual morphs in ruffs (10, 11).

The non-recombining portions of sex chromosomes are a special type of supergene because the two sexes are interdependent (12). The large body of research on the evolution of sex chromosomes constitutes a good starting point to understand supergene evolution. Evolutionary conflict over sexual phenotypes is believed to cause an enrichment of sexually-biased genes in the sex chromosomes, where the Y (or Z) variant is masculinized, and the X (or W) variant is feminized (13, 14). This pattern occurs in many established sex chromosome systems (15–18). Studies focusing on young sex chromosome systems, however, suggest that the role of evolutionary conflict in driving sex chromosome evolution has been overestimated (19–24). Instead, sex chromosome evolution could be driven mostly by processes unrelated to phenotypic differences between sexes (5, 25, 26). For example, gene-specific dosage compensation can be favored in response to mutations that change coding sequences (27) or expression levels (28, 29) in the non-recombining variant. In sum, processes derived from gene degeneration and selection for alternate genomic optima to overcome evolutionary conflict can play a role during the different stages of sex chromosome evolution (30).

We aim to understand the relative importance of evolutionary conflict and the consequences of recombination suppression in shaping the early evolution of a supergene. For this, we focus on the social supergene of the red fire ant *Solenopsis invicta*. Two social forms coexist in this species: colonies either have one or multiple queens. This social polymorphism is associated with multiple behavioral and physiological traits, all of which are controlled by a pair of “social chromosomes”, SB and Sb, that carry distinct supergene variants (31). In single-queen colonies, all workers and queens are SB/SB homozygotes. In multiple-queen colonies, all egg-laying queens are SB/Sb heterozygotes, but workers can either be homozygous or heterozygous. Sb/Sb queens are rare in the wild and their offspring are unviable (32, 33). Recombination is severely repressed between SB and Sb (31, 34) due to the presence of at least 3 inversions (35, 36). Similarly to a Y chromosome, Sb thus lacks recombination opportunities. The lack of recombination reduces the efficacy of selection in the Sb variant and is likely responsible for its ongoing degeneration through the accumulation of deleterious mutations (34, 35).

The extent to which a lack of recombination leads to gene depletion is variable. For example in Y chromosomes of Poeciliid fish, variation among taxa in gene depletion correlates with the intensity of sexual conflict (37). In the fire ant, the vast majority of genes are intact in both variants of the social chromosome supergene (31, 38). This lack of gene depletion is likely due to two main factors. First, the variants of the social chromosome only diverged recently (approximately 1M years ago (31, 38)). Additionally, male ants are haploid and therefore a haplotype with a deleted gene would be fully exposed to selection in males. The resulting selection against deletion would be much higher than in diploid species which would still carry an unaffected gene copy (35, 39).

Evolutionary conflict and the suppression of recombination should leave distinct signatures on patterns of gene expression. On one hand, conflict between single- and multiple-queen phenotypes should favor the accumulation of alleles adapted to the multiple-queen environment on the Sb haplotype. Under this scenario, we would expect that genes expressing the Sb more than the SB haplotype would tend to show an overall pattern of higher expression in multiple- rather than single-queen colonies. On the other hand, gene degeneration in the Sb haplotype should lead to lower expression of Sb alleles and perhaps gene-specific dosage compensation of corresponding SB alleles in dosage-sensitive genes. Here, we test these ideas by generating detailed genomic and transcriptomic data: we sequenced genomes of fire ants from their South American native range and combined these with existing genomes from the invasive North American range to identify genes with fixed differences between the SB and Sb variants of the social chromosome. To detect differences in expression between SB and Sb alleles, we performed RNA sequencing (RNA-seq) from SB/Sb individuals from the South American range of the species. We used three body parts of queens (head, thorax and abdomen) and whole bodies of workers. We then compared these expression patterns with those obtained from comparisons between social forms. Our results are consistent with evolutionary conflict shaping the evolution of the fire ant supergene in combination with a strong influence of the consequences of reduced recombination.

## Results

### Hundreds of genes have fixed allelic differences between supergene variants

To identify differences between supergene variants we obtained a total of 408× coverage of genome sequence from 20 haploid SB males and 20 haploid Sb males. Thirteen within each group (65%) are from the native South American range whereas the rest are from an invasive North American population (31). By comparing the two groups of males we identified 2,877 single nucleotide polymorphisms (SNPs) with one allele in all SB individuals and a different allele in all Sb individuals, affecting 352 genes (Supporting Information, Table S1). Among the 3.4% of SNPs affecting coding sequence, almost half change the amino-acid sequence (non-synonymous), with one change to a premature stop codon. The remaining SNPs were in intergenic (36.1%), intronic (58.0%) or untranslated regions (2.5%).

Because the invasive North American population went through a severe bottleneck in the 20th century (40), we repeated the analysis after separating populations. We found 252 additional SNPs with fixed differences between SB and Sb individuals in South America, and an additional 23,022 fixed differences between SB and Sb in North America. The latter number is 4-fold higher than expected due to differences in sampling size alone and is in line with lower genetic diversity of both supergene variants in North America due to the invasion bottleneck (40).

### Seven genes have consistent variant-specific expression patterns in all populations

Because most coding sequences are intact in both variants of the supergene, we asked whether variant-specific allelic expression biases may occur. We generated individual-specific RNA-seq data from whole bodies of SB/Sb workers and from abdomens, thoraces, and heads of SB/Sb virgin queens collected in South America. To add additional robustness to our analysis, we additionally incorporated existing RNA-seq gene expression data from pools of whole bodies of SB/Sb queens collected in the USA and Taiwan (41, 42).

Among the 352 genes with fixed differences between SB and Sb, 122 had sufficient expression for analysis of differences between alleles. Using a linear mixed-effects model, we found that seven of the genes had consistent expression differences between variants across all populations (Benjamini-Hochberg (BH) adjusted p<0.05, Figure 1, Supporting Information, Table S2). Expression bias went in both directions: the Sb variants of “pheromone-binding protein Gp-9/OBP3” (LOC105194481), “retinol-binding protein pinta-like” (LOC105199327) and LOC105193135 (uncharacterized) were consistently more highly expressed, while the SB variants of “ejaculatory bulb-specific protein 3” (LOC105199531), “carbohydrate sulfotransferase 11-like” (LOC105193134), “calcium-independent phospholipase A2-gamma” (LOC105203065) and LOC105199756 (uncharacterized) were consistently more highly expressed.

**Figure 1.**
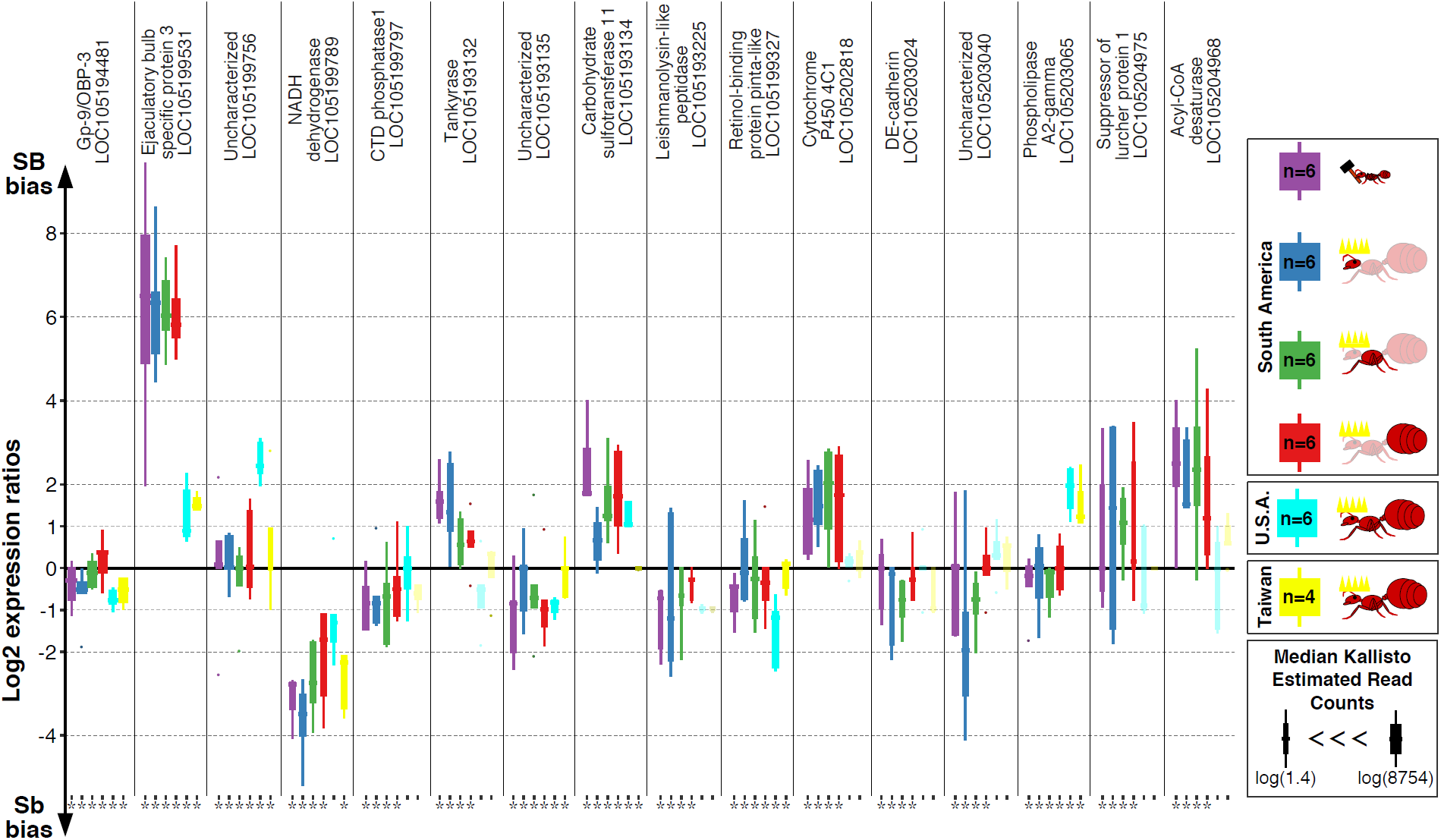
Allele-specific expression for genes in the fire ant social supergene in heterozygous SB/Sb individuals. Differences in expression (y axis) between the variants of the social chromosome in whole bodies of South American workers (purple), heads (blue), thoraces (green) and abdomens (red) of South American queens, whole bodies of North American queens (light blue) and whole bodies of Taiwanese queens (yellow). We show relative expression levels for the 16 genes from the social chromosome supergene region which have significant allele-specific expression differences between variants in at least one population. Significance for each gene for each comparison is indicated by an asterisk at the bottom of each plot, and non-significant comparisons are semi-transparent (Benjamini-Hochberg (BH) adjusted p<0.05 from the joint linear mixed-effects for Taiwanese, South and North American populations; BH adjusted p<0.05 from DESeq2 Wald tests for analyses restricted to South America or Taiwan). Each box shows the distribution of logarithm 2 expression ratios between SB and Sb per caste/body part/population. Genes with values above the solid line (Log2 expression ratios=0) have higher SB expression; those with values below the line have higher Sb expression. A log2 expression ratio greater than 1 or smaller than -1 represents a two-fold gene expression difference in either direction. Genes are in chromosomal order.

We additionally identified differences between expression of SB and Sb-linked alleles using a population-specific approach for South American and also independently in North American and Taiwanese populations. We found 124 expressed genes containing fixed differences between variants in the native South American range, 343 and 374 such genes in the invasive populations of North America and Taiwan respectively. Most of the genes with sufficient expression data and fixed differences between variants in South America (122 out of 124, 98.4%) could also be examined in the North American and Taiwanese datasets (Figure 1-Figure supplement 1).

We applied DESeq2 (v1.14.1 (43)) to each population to detect allelic expression differences. Within the South American data we generated, this approach detected 12 genes with significant allele-specific expression, 6 being more highly expressed in SB and 6 in Sb (Figure 1-Figure supplement 2, Table S3a). Of these, three had been detected by the linear mixed effects model using all populations (“carbohydrate sulfotransferase 11-like”, “ejaculatory bulb-specific protein 3” and “uncharacterized LOC105193135”). We found no general bias in expression towards either variant (median of log2 expression ratios=-0.07; Wilcoxon sum rank test p=0.27).

We repeated this analysis using gene expression data from pools of SB/Sb queens from North American populations of *S. invicta* (41) (Figure 1-Figure supplement 3, Table S3b). Twenty nine of the 343 genes showed significant allele-specific expression. Of these, six had been detected by the linear mixed effects model using all populations. There was no directional bias, with 16 of the genes being more highly expressed in SB, and 13 in Sb (binomial test p=0.7; median of log2 expression ratios=6×10^−17^; Wilcoxon sum rank test p=0.40).

Finally, we also performed this analysis using pools of SB/Sb queens from Taiwanese populations (42) (Figure 1-Figure supplement 4, Table S3c). Out of the 374 genes analyzed, 19 showed significant allele-specific expression. Of these, three had been detected in the joint linear mixed effects model. Additionally, 12 of the genes with significant allele-expression differences between variants found in Taiwanese populations were independently found using North American populations too. Again, we found no directional bias in allele-specific expression across the supergene (8 genes more highly expressed in Sb, and 11 in SB; binomial test p=0.64; median of log2 expression ratios=-0.02, Wilcoxon sum rank test p=0.11).

These two additional analyses are thus consistent with the findings from the overall model reported above, and also uncovered population-specific expression.

### Supergene allele expression differs based on population ancestry but not body parts

If allele-specific expression patterns of genes within the supergene are mostly driven by selection for social form-specific functions, we would expect allelic expression biases to differ between tissues. The South American RNA-seq data we generated provided the opportunity to test this idea because it includes multiple body parts and castes (head, thorax and abdomen in queens, and whole body in workers). We found no significant effect of body part or caste on allelic expression for any gene (DESeq2’s Logarithmic Ratio Test and all pairwise Wald comparisons between interaction terms; all BH adjusted p>0.05). This finding suggests that allelic expression within the supergene is not fine-tuned for specific functions in different tissues. We found additional support for this finding by detecting allelic expression differences between body parts for genes in normally recombining parts of the genome, despite having less power to do so than in the supergene region (see Methods).

The progressive degeneration of the Sb variant of the supergene through the accumulation of deleterious mutations (35) could affect gene expression. Under this scenario, we expect allelic expression patterns within the supergene to differ more between populations that are more distantly related. Our inclusion of three populations with different levels of relatedness provided the opportunity to test this hypothesis. For each pair of populations, we calculated correlations of the log2 ratios between SB- and Sb- allelic expression (Figure 1-Figure supplement 5). The log2 expression ratios between North American and Taiwanese populations were strongly correlated (Spearman’s r^2^=0.67), likely because the North American and Taiwanese populations are highly related (44). In contrast, the correlation between either invasive and the South American population was weak (Spearman’s r^2^=0.21 and 0.18 for the South America-North America and South America-Taiwan correlations respectively). This result was further supported by an additional linear mixed-effects model incorporating all data. This model shows that ancestry of the populations explains more of the variance observed in allelic expression than the continent on which they are (interaction between North American-Taiwanese populations and gene effect in predicting log2 expression ratios: F=3.42, p<10^−15^; interaction South-North American populations and gene effect in predicting log2 expression ratios: F=0.94, p=0.65).

These results support the idea that genomic architecture plays a major role in defining supergene allelic expression.

### Overrepresentation of socially biased genes in the social supergene

Differential selection for single-queen and multiple-queen colonies leads to the prediction that genes with socially biased expression should be overrepresented in the supergene region. To test whether this occurs, we compared gene expression between egg-laying queens from single-queen and from multiple-queen colonies. We identified 293 such socially biased genes for which chromosomal locations are known (Supporting Information, Table S4). Such genes were indeed overrepresented in the supergene region (Figure 2a, 33 out of 293, 12 expected by chance, χ^2^=29.7, p<10^−7^). Additionally, the vast majority of socially biased genes (274 out of 293, *i.e*., 94%) were more highly expressed in multiple-queen colonies than in single-queen colonies (binomial test, p<10^−7^, Figure 2b). We independently confirmed this result by comparing expression profiles of queens from multiple-queen colonies from another RNA-seq dataset (41) to those of queens from single queen colonies (45). For this, the samples were normalized together using the default DESeq2 method. Because these samples belong to different datasets, there are too many confounding factors to estimate significance levels of differential expression for each gene. We thus used a different approach. We first performed a simple DESeq2 analysis to obtain the log2 expression ratios of this comparison. We then compared the log2 expression ratios using a linear regression to those obtained from the comparison using only the Morandin *et al.* (45) data to detect patterns shared by the two datasets. The strong trend we found (49% of the variance in log2 expression ratios explained by social form ± a standard deviation of 2.9%, p<10^−16^)) confirmed previous results.

**Figure 2.**
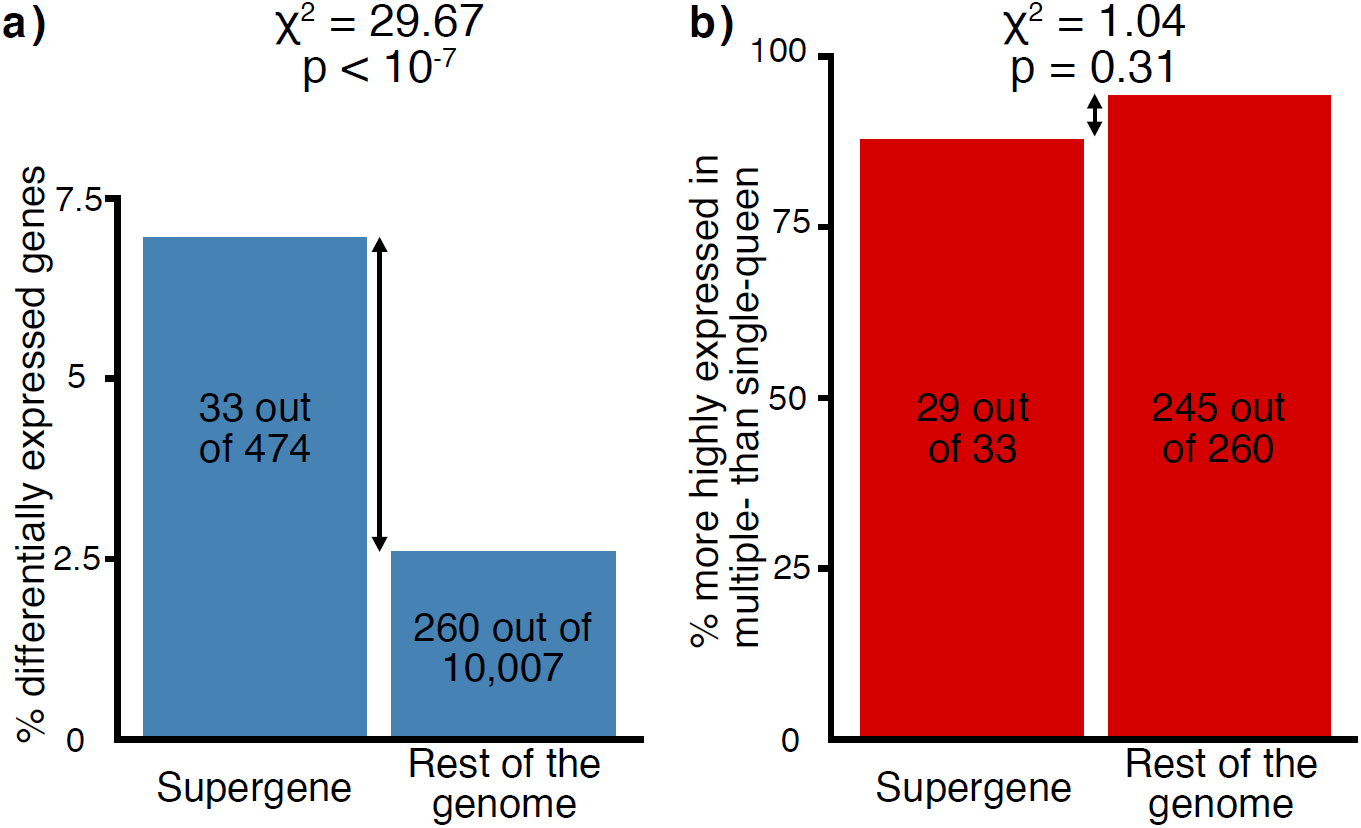
Distribution of socially biased genes in the genome of the red fire ant within (left bars) and outside (right bars) the supergene region **a)** for genes with significant expression differences between social forms and **b)** for socially biased genes that are more highly expressed in multiple-queen colonies than in single-queen colonies. The vast majority of genes with differential expression between social forms are more highly expressed in multiple-queen colonies. The proportion of genes that are more highly expressed in multiple-queen colonies is similar within and outside the supergene, which shows that expression bias towards multiple-queen colonies is not a unique feature of the supergene.

### Genes with higher expression of Sb than SB alleles are socially biased towards higher expression in multiple-queen colonies

Similarly to the Y chromosomes in males, because Sb is present only in multiple-queen colonies, it should accumulate alleles beneficial for multiple-queen colonies. When a gene has sexually-biased gene expression, this is generally interpreted as a sign that this expression is beneficial to that sex (47). If we assume that socially biased expression similarly indicates a benefit to the particular social form, genes with higher expression of Sb than SB alleles should be more highly expressed in multiple-queen than in single-queen colonies.

Information regarding gene expression differences between social forms exists only for North American populations. To match genes with differences in expression between social forms with genes showing allele-specific expression differences between supergene variants, we analyzed data from North American populations only. Among the genes in the supergene region, 294 had sufficient data for both comparisons (Figure 3 and Figure 3-Figure supplement 1). Most of these genes (256; 87%) had no significant differential expression patterns (Figure 3a) whereas eight genes (3%) had both allele-biased and socially-biased expression. Their expression was strongly directionally biased, with the majority of these genes having higher expression in Sb and in multiple-queen colonies (5 out 8 Sb biased genes were more highly expressed in multiple-queen colonies (Figure 3d); compared to 1 out of 15 for SB biased genes, χ^2^=5.8, p=0.02). We additionally tested whether this was a general pattern for the supergene region. To do this, we extended the analysis to search for the same trends using all 294 genes in the supergene region. With this data we fitted a linear model estimating changes in allele-specific expression based on gene expression differences between social forms (Fig 4). The model showed that higher expression of the Sb allele results in higher expression in individuals from multiple-queen than single-queen colonies. This approach additionally identified a trend that went undetected when considering individual genes: higher expression of the SB alleles in heterozygous individuals generally corresponds to lower expression in multiple-queen than in single-queen colonies. We find that compared with the standard linear regression (straight black line in Figure 4), there is a slightly better fit for all values for the linear regression of allelic expression bias (*P*_*B*_) on the gene expression bias towards multiple-queen colonies (*P*_*MQ*_; whereby *P*_*B*_*=(1 - P*_*MQ*_) / *P*_*MQ*_; curved red line in Figure 4). For both relationships, the significant increase in *P*_*B*_ for genes with relatively low expression in multiple-queen colonies (*P*_*MQ*_<0.5; p<10^−5^) suggests that that expression of the Sb allelic variant is depleted at these loci. We additionally show that this pattern is very unlikely to be explained solely due to Sb degeneration, consistent with an adaptive role of some of the genes with Sb biased expression. A model explaining the observed expression patterns based on Sb degeneration alone (*i.e.*, based on allelic bias alone: P_*MQ*_ = 1 / (2*P*_*B*_ + 1); purple line in Figure 4) poorly predicts the data and is significantly different from the best fit model (p<10^−5^).

**Figure 3.**
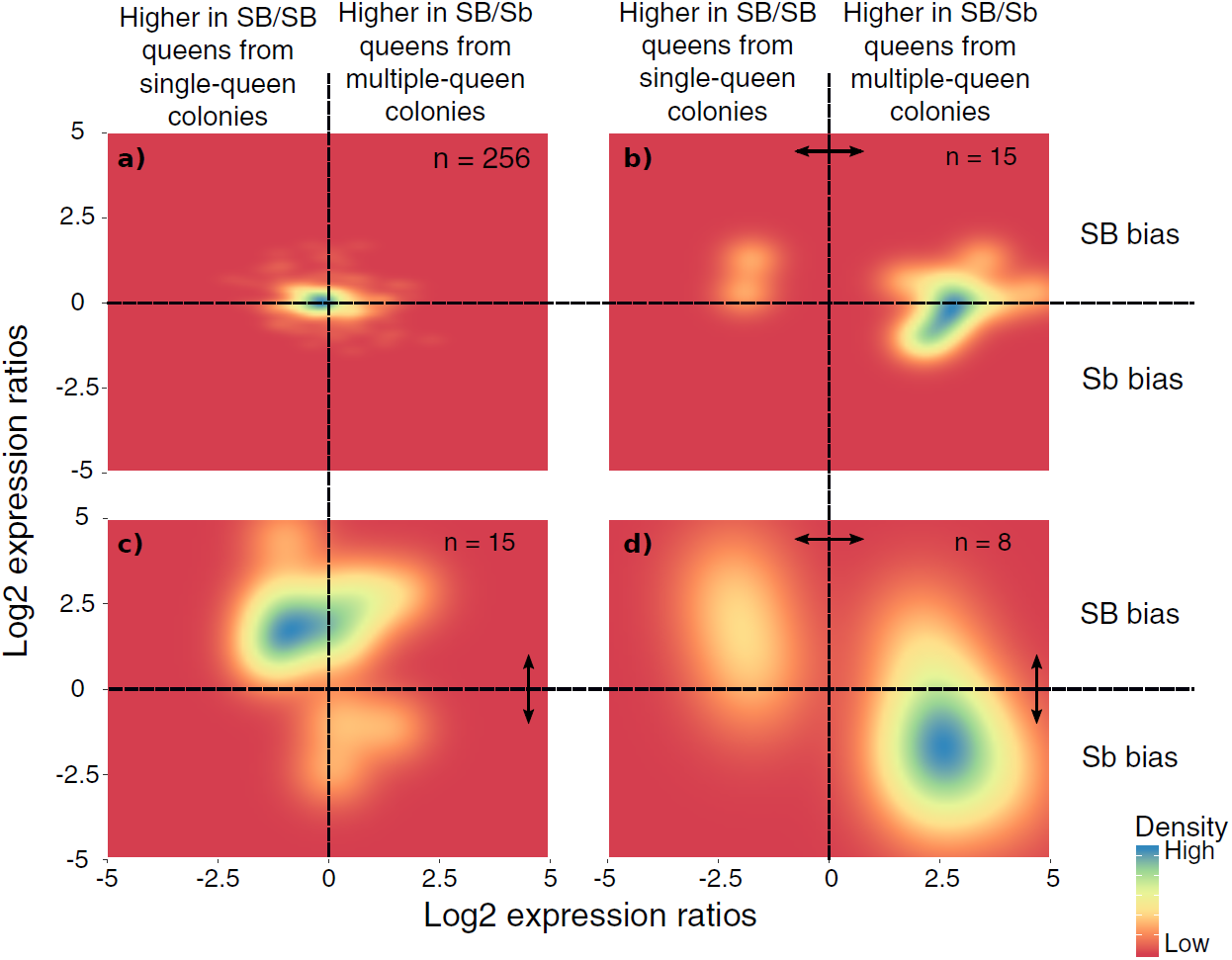
Distribution of differences in gene expression between social forms and between supergene alleles. X axes indicate ratios of expression between SB/Sb queens and SB/SB queens. Y axes indicate allelic expression ratios in SB/ Sb queens. Both ratios use a log2 scale whereby log2=0 indicates absence of differences. The colors are proportional to the number of genes with the focal expression. Double-headed arrows indicate significant expression differences. Panel **a)** expression ratios for genes showing no difference in either comparison. The remaining three panels summarize the expression patterns for: **b)** genes with significant expression differences between SB/Sb and SB/SB queens only – these are biased towards higher expression in multiple-queen colonies (binomial test, p=0.007); **c)** genes with significant expression differences only between SB and Sb alleles within SB/Sb individuals – these are biased towards higher expression in the SB variant, in line with a dosage compensation mechanism (binomial test, p=0.03); **d)** genes with significant expression differences between SB/Sb and SB/SB queens and between the SB and Sb variants in SB/Sb queens – the genes with higher expression of the Sb allele (x>0) tend to be more highly expressed in queens from multiple-queen colonies (y<0), in line with evolutionary conflict between social forms (5 out of 8 Sb biased genes with bias towards multiple-queen colonies, compared with 1 out of 15 for SB biased genes χ^2^=5.8, p=0.02).

**Figure 4.**
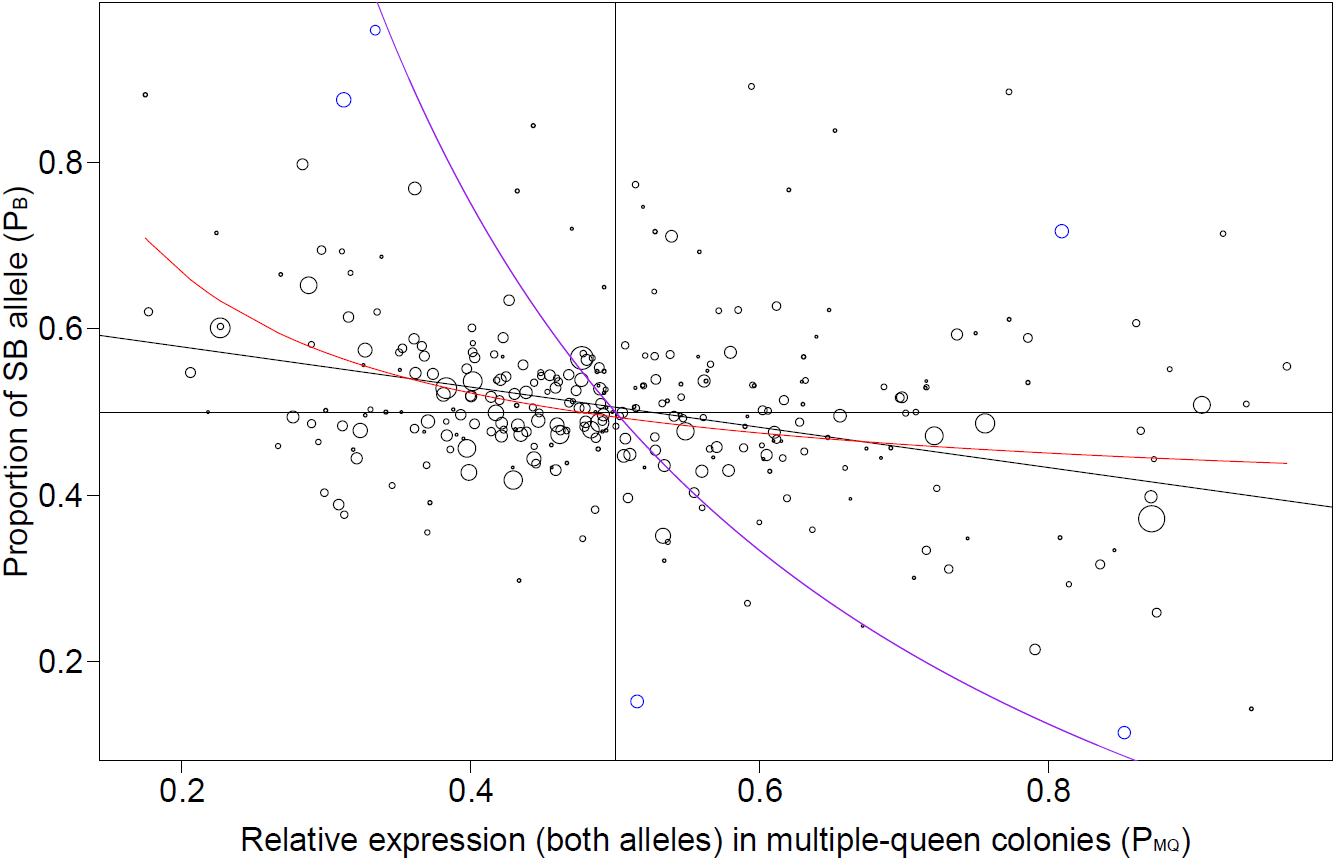
Relationship between expression bias of the B allele compared to the b allele in SBSb individuals (P_B_) and gene expression bias of multiple-queen colonies compared to single-queen colonies (P_MQ_). Each point is one of 294 genes within the supergene (North American data). Point size is proportional to the mean expression in SBSb individuals. The values were calculated as *P*_*B*_ = *x*_*B*_/(*x*_*B+*_*x*_*b*_) and *P*_*MQ*_ = *x*_*MQ*_/(*x*_*SQ+*_*x*_*MQ*_); where *x*_*B*_ and *x*_*b*_ are the expression levels of SB and Sb alleles, and *x*_*SQ*_ and *x*_*MQ*_ are the expression levels in single-queen and multiple-queen colonies. Values below 0.5 in the x axis therefore indicate higher expression in SB/SB queens from single-queen colonies, and values above 0.5 indicate higher expression in SB/Sb queens from multiple-queen colonies. Values above 0.5 in the y axis indicate bias towards the expression of the SB allelic variant; below 0.5 indicate bias towards the Sb variant. The straight black line is the linear regression, the curved red line is the regression on allelic bias as a function of social expression bias (1-P_MQ_/P_MQ_). Both relationships are highly significant (p<10^−5^). The purple line shows the predicted fit if the pattern of expression was due to Sb degeneration alone, *i.e.*, based only on allelic differences between variants (1/ (2P_B_+1)). This model is a poor predictor of the data and is significantly different from the best fit model (p<10^−5^). The observed enrichment of multiple-queen genes in Sb is therefore very unlikely due to Sb degeneration alone.

### Genes with no social bias tend to have allele-specific expression biased towards the SB variant

If purifying selection is relaxed for the majority of genes in the Sb variant of the supergene, they should be accumulating deleterious mutations. In consequence, the expression of Sb alleles for such genes would decrease or be downregulated. If stoichiometric ratios require such genes to be expressed at specific levels, we would expect their “healthy” SB alleles to compensate for “faulty” Sb variants. In such cases of gene-specific dosage compensation we would thus expect allelic bias towards higher expression of SB alleles, but no gene expression differences between social forms.

Fifteen of the 294 genes in the previous section had allele-specific expression differences but no differences in gene expression between social forms (5%; Figure 3c). Most of these genes (12) had higher expression in SB, with only 3 being more highly expressed in Sb (binomial test, p=0.03). In line with this, the median SB-Sb expression ratio across all 15 genes was 5.9 (Wilcoxon signed-rank test p=0.0008 for differences against a ratio of 1). The bias in expression towards the SB alleles of genes with no expression differences between social forms is consistent with gene-specific dosage compensation taking place in the supergene region.

## Discussion

In the fire ant, a supergene system with two variants (SB and Sb) controls whether colonies have one or multiple queens. We compared gene expression patterns between the SB and Sb variants of the social supergene within heterozygote SB/Sb individuals which exist only in multiple-queen colonies, and between queens from single-queen and multiple-queen colonies. Our results show patterns of expression consistent with evolutionary conflict and with degradation of Sb. Furthermore, our work highlights candidate genes potentially responsible for differences between social forms.

### A balance between Sb degeneration, dosage compensation, and selection for alternate social forms

We found that in each population, 5% to 10% of the genes in the supergene region showed allele-specific expression differences between the SB and Sb variants. Seven of these genes show consistent SB-Sb allelic expression biases across all the populations. It is tempting to conclude that gene expression differences arose through selection, as a consequence of evolutionary conflict between the single-queen and multiple-queen phenotypes. Our other results, however, suggest that first level interpretations of expression patterns can be too simplistic, as they ignore the impacts of supergene degeneration on gene expression.

We found enrichment of genes with expression differences between single-queen and multiple-queen colonies in the supergene region (Figure 2). Two alternative hypotheses can explain this enrichment. The first hypothesis is that evolutionary conflict drives differences in gene expression between social forms in a similar way to the generally accepted model of sex chromosome evolution (30). Assuming that high expression levels reflect a beneficial effect for the bearer (14), evolutionary conflict between social forms should lead to increased expression of genes favoring each of the social forms in each of the supergene variants (1, 48). The alternative hypothesis is that the differences in expression between social forms are non-adaptive, instead reflecting genetic differentiation between SB and Sb at gene regulatory sites in the supergene region. Although some of this differentiation could be neutral, most is expected to be deleterious, as a result of the low efficacy of purifying selection in Sb caused by absence of recombination (*i.e.*, Hill-Robertson interference) (34). Indeed, point mutations (49, 50) and insertions of transposable elements (23) can cause changes in gene expression, and Sb is enriched both types of mutations (34, 51).

We performed a joint analysis of gene expression differences between social forms and between supergene variants. For this analysis, we used only allele-specific information from North American colonies because this was the only population for which we had information for social form differences. We found that 23 genes in the supergene had fixed differences between variants and differential expression between the social forms. For fifteen of these genes, SB and Sb alleles had similar expression levels. Five of the eight genes with differences in expression between SB and Sb alleles followed a single pattern: higher expression in the multiple-queen social form, and higher expression of Sb than SB alleles in SB/Sb heterozygotes. Assuming that higher expression in one of the social forms indicates adaptive value, we conclude that the Sb alleles of these genes likely evolved as a response to selection for adaptation to the multiple-queen social phenotype.

We additionally observed a depletion of multiple-queen biased loci in the SB variant. Similarly to what we observed for Sb, this pattern could be interpreted as selection against the expression of single-queen biased loci in SB. However, the pattern of enrichment in Sb is stronger than the depletion in SB, and is stronger than what would be expected due to Sb degeneration alone. The different strengths of these patterns may reflect an important characteristic of the social chromosome system: that the Sb supergene variant only occurs in multiple-queen colonies, whereas the SB variant occurs both in single-queen colonies and in multiple-queen colonies. This characteristic of the system raises the expectation that selection is effective at accumulating alleles benefitting multiple-queen colonies in Sb, but less effective at accumulating alleles benefitting single-queen colonies in SB, which is consistent with the expression patterns we observed. Similarly, in some XY systems, the X chromosome lacks female-biased alleles (52).

Interestingly, the supergene also included 15 genes with allelic expression biases but no difference between social forms (Figure 3c, Fig S5). For the majority of genes with this pattern, the allele carried by SB had higher expression than its Sb counterpart. This result is consistent with a gene-specific dosage compensation mechanism to ensure stable overall expression despite decreased expression of the Sb variant. Two processes could explain why dosage compensation might have arisen for these genes. First, selection could favor lower expression of the Sb allele if random mutations lead to mis-expression, or to the production of non-functional proteins. Alternatively, mutations in regulatory elements in the Sb variant could have directly decreased expression levels of Sb alleles. In both cases, higher expression of the “healthy” SB variants would then have been selected for to offset the lower expression of Sb. Regardless of what is driving dosage compensation in this system, the overall pattern is similar to reports in other young supergene systems (20, 21, 53), where compensation occurs for individual genes rather than globally, as seen in, for instance, some mammalian X chromosomes (54). Furthermore, these two hypotheses underlying potential causes for dosage compensation could act simultaneously. Indeed, most of the genes with low Sb expression (92%, 11 out of 12) had more than one mutation in potentially regulatory upstream regions and slightly more than half (58%, 7 out of 12) were affected by nonsynonymous coding substitutions. This proportion of nonsynonymous mutations is similar to that observed across the supergene, where roughly half of the fixed exonic mutations between SB and Sb impact the protein sequence (47.7% nonsynonymous, 52.3% synonymous). This finding is consistent with ongoing gene degeneration, in line with previous results in this (34) and other young supergenes (9).

Gene degeneration could also contribute to the relatively small number of genes with SB and multiple-queen biased expression. One possibility is that gene degeneration leads to low expression of the Sb allele, which has two effects: a relatively low expression of the affected gene, combined with bias towards the expression of the SB allele (rather than Sb). A second interpretation is that the low expression of a gene in multiple-queen colonies is indicative of relaxed selection on that gene. Relaxed selection would in turn facilitate degeneration of the Sb allele, and its consequent reduction in expression. These explanations are not mutually exclusive.

Remarkably, the degree of differential allelic expression of SB and Sb alleles was consistent across body parts for all the loci where allelic bias was detected (Figure 1). This pattern contrasts with human data, in which tissue-specific selection resulted in allele-specific expression variation across tissues (43). Similarly, in the fire ant we might have expected that selection would aim to fine-tune gene expression independently in each tissue to accommodate for tissue-specific functions in each social form. The young age of the supergene (31) in combination with low selection efficacy may have precluded such fine-tuning. As a result, strong selection for a particular level of allele-specific expression in one body part (*e.g.*, in antennae), could result in a consistent allele-specific expression pattern across tissues, even if this has mildly deleterious effects (*e.g.*, in the gut).

The absence of allelic differences between body parts and castes contrasted with our finding strong effects of population of origin on allelic expression: gene expression differences between SB and Sb were strongly correlated between North American and Taiwanese populations, despite the data from the two populations being from different studies. The North American population derived from the South American population over the last century, and the Taiwanese population derived from the North American population within the last 30 years (44). Both populations thus have lower genetic diversity overall, and in the supergene region in particular (34, 44). In line with this finding, the correlation in gene expression patterns within the supergene was weak between South American and Taiwanese or North American populations. This strong effect of ancestry suggests an important role of genomic architecture in defining expression patterns within the supergene.

Overall, our results show that gene degeneration plays a major role in shaping the expression patterns within the supergene. Indeed, the fixed differences in allele-specific expression across populations that we report may be consequences of recombination suppression rather than adaptations arising from evolutionary conflict. To distinguish between these two processes, it is relevant to put allele-specific biased expression in the context of gene expression differences between social forms. By doing so, we highlight several candidate genes that could play key roles in defining social forms.

### Candidate genes for differences between social forms

The approaches underpinning our analyses are unable to detect allelic differences in genes absent from the reference genome such as OBP-Z5, a putative Odorant Binding Protein exclusive to Sb (38). However, our analysis does single out candidate genes potentially contributing to the social polymorphism of the fire ant (lists of all genetic differences between SB and Sb are in Table S1; lists of genes that showed expression differences in any comparison are in Tables S2, S3 and S4). Three genes in particular stood out because they were differentially expressed between social forms and also had variant-specific allele expression in all populations. For the first gene, “Pheromone-binding protein Gp-9” (LOC105194481), also known as OBP-3, the Sb allele was more highly expressed. For decades, this gene has been a candidate effector for social form differences (34, 55), yet its linkage to hundreds of other genes in the supergene led to doubts that its association to social form is any more than coincidental. We found five fixed differences between SB and Sb for this gene, four of which could have an impact on protein efficiency (consistent with findings from Krieger and Ross (56)). For the second gene, “Ejaculatory bulb-specific protein 3” (LOC105199531), which also contains an insect odorant binding protein domain (InterPro IPR005055), the SB allele was more highly expressed. Orthologs of this gene are associated with mating (57) in *Drosophila melanogaster*, sexual behavior in a moth (58), subcaste differences in bumblebees (59), venom production in social hornets (60) and caste differences in the termite *Reticulitermes flavipes* (61). Finally, LOC105199327 is similar in sequence to Pinta retinol-binding proteins, which are linked to pigment transport and vision in *D. melanogaster* and the butterfly *Papilio xuthus* (62). In sum, all these candidate genes have putative functions related to environmental perception, in line with the complex social phenotype requiring subtle changes in environmental perception or signaling.

## Conclusions

By analyzing allele-specific expression patterns between the SB and Sb variants of the fire ant supergene we found consistent differences across three populations. Such strong patterns can naively be assumed to be indicative of adaptive processes emerging from evolutionary conflict between social forms. We show, however, that the evolutionary forces shaping expression patterns in the supergene are complex and must be interpreted with care. In particular, genes with higher expression of the SB than the Sb allele tend to either lack expression differences between social forms or have lower expression in multiple-queen than in single-queen colonies. Both patterns are consistent with the idea that recombination suppression leads to degeneration in Sb and thus lower expression of Sb alleles for some genes. In some cases, a dosage compensation mechanism through higher expression of the healthy SB allele leads to similar expression levels in both social forms. In cases where no dosage compensation occurs, overall expression is lower in multiple-queen colonies than in single-queen colonies.

Conversely, we also show that genes with higher expression of the Sb than the SB allele also tend to have expression biased towards multiple-queen colonies. This pattern is consistent with evolutionary conflict favoring the accumulation of beneficial alleles for multiple-queen individuals in Sb, the supergene variant found only in this social form.

Our study shows that multiple complex evolutionary forces can simultaneously act on a young supergene system. In particular, it highlights that allele-specific expression patterns alone are insufficient for inferring whether they are adaptive, deleterious or phenotypically neutral. Instead, putting such expression differences into broader contexts is needed to draw reasonable conclusions. By applying this idea to our data we highlight genes which have molecular roles that could affect perception or signalling of the social environment.

## Materials and Methods

### RNA sequencing of fire ants

We used three previously published and generated one novel fire ant RNA-seq datasets. Wurm *et al* (41) obtained whole-body RNA-seq data from six pools of 4 egg-laying SB/Sb queens, each from a multiple-queen colony from Georgia, USA. Morandin *et al* (45) obtained whole-body RNA-seq data from 6 samples, each being a pool of 3 queens from a single-queen or a multiple-queen colony (3 replicates per social form) from Texas, USA. Due to low mapping quality, we eliminated one queen sample from one of the multiple-queen colonies. All the queens were mature and egg-laying (C. Morandin, personal communication), thus queens from multiple-queen colonies carried the SB/Sb genotype (Keller and Ross 1998). Additionally, we manually checked the resulting RNA reads for heterozygous positions at key supergene markers using the IGV gene browser v2.4.19 (63). Fontana *et al* (42) generated RNA-seq data from 4 samples of SB/Sb queens from multiple-queen colonies from Taiwan. Each sample is a pool of whole bodies from two virgin queens (more details in Supporting Information, Table S5a).

All three published datasets are from pools of whole bodies from the invasive ranges of *S. invicta.* The Taiwanese invasive population of red fire ants is derived from that of North America (44). Both invasive populations will therefore be considered as a single distinct from that of South America, for the purposes of this study. Because comparisons of whole bodies can be confounded by allometric differences (64) and genetic diversity is reduced among Sb haplotypes in the invasive populations of North America (34), we generated a new gene expression dataset. From the native South American range of this species, we collected 6 multiple-queen colonies (collection and exportation permit numbers 007/15, 282/2016, 433/02101-0014449-4 and 25253/16). We confirmed the social form of each colony using the Krieger and Ross assay (56) on a pool of DNA from 10 randomly chosen workers. Colonies were kept for 6 weeks under semi-controlled conditions (natural light, room temperature, cricket, mealworm and honey water diet) before sampling. From each colony, we snap-froze (between 12:00 and 15:00 local time) one worker and one unmated queen for gene expression analysis. To partly control for allometric differences between genotypes, we separated each queen into head, thorax and abdomen. This was done in petri dishes over dry ice using bleached tweezers. In total, we had 24 samples for RNA extraction: six whole bodies of workers and six replicates of three body parts from queens (more details in Supporting Information, Table S5b).

We extracted RNA and DNA from each sample using a dual DNA/RNA Tri Reagent based protocol (Supplementary file 1). We applied the Krieger and Ross assay (56) on the extracted DNA to identify only individuals with the SB/Sb genotype. Once RNA was extracted, we prepared Illumina sequencing libraries from total RNA using half volumes of the NEBNext Ultra II RNA Library Prep Kit. Raw RNA and the libraries were quality checked on an Agilent Tapestation 2200; library insert size averaged 350 bp. An equimolar pool of the 24 libraries was sequenced on a single lane of Illumina HiSeq 4000 using 150 bp paired-end reads. This produced an average of 14,848,226 read pairs per sample (maximum: 27,766,980; minimum: 6,015,662. Raw RNA-seq reads for all samples are on NCBI SRA (PRJNA542606).

For all datasets, we assessed read quality using fastQC (v0.11.5; http://www.bioinformatics.babraham.ac.uk/projects/fastqc/). Raw reads for all samples were of sufficient quality to be used in subsequent analysis. We removed low quality bases using fqtrim with default parameters (v0.9.5; http://ccb.jhu.edu/software/fqtrim/), and Illumina adapters using Cutadapt v1.13 (65). We then generated a STAR v2.5.3a (66) index of the *S. invicta* reference genome (version gnG; RefSeq GCF_000188075.1 (41) while providing geneset v000188075.1 in GFF format through the “sjdbGTFtagExonParentTranscript=Parent” option. As recommended by the developers of STAR, we aligned each sample to the reference twice, using the “out.tab” file for the second run, and set “sjdbOverhang” to the maximum trimmed read length minus one, *i.e*., 74 for the Wurm *et al.* data (41) and Morandin *et al.* data (45), 125 for the Fontana *et al.* data (42) and 149 for the South American data we generated here. Alignments were run using GNU Parallel v20150922 (67). All steps and downstream analyses were performed on the Queen Mary University of London’s Apocrita High Performance Computing Cluster (68).

We further assessed aligned reads (*i.e.*, BAM files) using MultiQC v1.5 (69) and the BodyGene_coverage.py script of the RSeQC toolkit v2.6.4 (70). We removed one sample from multiple-queen colonies in the Morandin *et al.* data from subsequent analyses due to poor alignment quality. None of the other BAM files showed markers of technical artefacts that could bias our results.

### Identifying SNPs with fixed differences between SB and Sb males

To detect allele-specific differences between SB and Sb we first identified SNPs with fixed differences between the SB and Sb variants. Because the patterns of genetic diversity differ between the invasive and South American *S. invicta* populations (71, 72), we estimated allele specific expression differences in the social chromosome independently for each population. For this we used haploid male ants because they can provide unambiguous genotypes. For the invasive populations, we identified fixed allelic differences between a group of 7 SB males and a group of 7 Sb males from North America (NCBI SRP017317) (31).

For the South American population, we sequenced the genomes of 13 SB males and 13 Sb males sampled from across Argentina. For each individual, we extracted 1 µg of genomic DNA using a phenol-chloroform protocol. The extracted material was sheared to 350 bp fragments using a Covaris (M220). We constructed individually barcoded libraries using the Illumina TruSeq PCR-free kit. The libraries were quantified through qPCR (NEB library quant kit). An equimolar pool of the 46 libraries was sequenced on a HiSeq4000 at 150 bp paired reads. This produced an average of 17,790,416 pairs of reads per sample, with a maximum of 38,823,285 and a minimum of 7,910,042. (Table S6, genomic reads of all samples deposited on NCBI SRA (PRJNA542606)). For each dataset, we identified fixed allelic differences between the group of SB males and the group of Sb males. We first aligned the reads of each sample to the *S. invicta* reference genome ((41); gnG assembly; RefSeq GCF_000188075.1) using Bowtie2 v2.3.4 (73). We then used Freebayes v1.1.0 (74) to call variants across all individuals (parameters: ploidy = 1, min-alternate-count = 1, min-alternate-fraction = 0.2). We used BCFtools (75) and VariantAnnotation (76) to only retain variant sites with single nucleotide polymorphisms (SNPs), with quality value Q greater than or equal to 25, and where all individuals had a minimum coverage of 1. To avoid considering SNPs erroneously called from repetitive regions that are collapsed in the reference genome, we discarded any SNP with mean coverage greater than 16 for the North American samples or 12 for the South American samples or where any individuals had less than 60% reads supporting the called allele. This last filtering step also acts to remove SNPs called from reads with sequencing errors. We then extracted only the SNPs located within the supergene (based on the genomic locations from Pracana *et al.* (34)) and with fixed differences between SB and Sb. This step was performed independently for each population. The two resulting variant call files were inspected using VCFtools v0.1.15 (77) and we manually ensured that all variants had the SB allele as reference and Sb allele as alternative. To test the effect of sample size differences between populations we downsampled the South American dataset to 7 pairs of SB and Sb males, matching the sample size in the North American dataset.

We extracted SNPs shared between South and North American populations using BCFtools isec v1.9 (75). We then used SNPeff (78) to characterize the effects of individual SNPs.

### Estimating read counts from alternate social chromosome variants in heterozygous individuals

Because the reference genome for *S. invicta* is based on an SB individual, read mapping could be biased towards the SB variant in heterozygous individuals, resulting in apparent overall expression of the reference variant (79). To overcome this potential artifact, we performed two alignments using STAR (with the same parameters as described in the “Generation of RNA-seq data” section): one with the regular SB reference genome and another with a modified reference genome in which we had replaced the fixed positions between variants in the supergene by those found in Sb. We then used the last steps from the WASP pipeline (80) to generate reference-bias free alignment files. We obtained allele-specific counts using GATK’s “ASEReadCounter” v 3.6-0-g89b7209 (81) using the fixed SNP differences between Sb and SB. To generate allele-specific read counts in a manner that eliminates potential reference bias, we first generated modified red fire ant reference genomes in which all the positions in the supergene had been changed to match the Sb alleles. We called BCFtools consensus v1.9 (75) once using North American Sb males and once using South American Sb males. We then aligned the RNAseq reads from each sample to the regular reference genome (version gnG; RefSeq GCF_000188075.1; (2)) and also, independently, to the modified references. For the reads from the Taiwanese population, we used the Sb reference using North American SNPs. For the alignment we used STAR with the same parameters as described above. We merged the two resulting BAM files from each sample using SAMtools v1.9 (75). The vast majority of reads would have been mapped to both references, and thus appear twice in the merged aligned file. To remove these duplicates in an unbiased manner, we used the “rmdup” script from the WASP pipeline (80). The resulting BAM files can be considered reference bias free alignments.

We added a reading group ID to each reference-bias free BAM file using the “AddOrReplaceReadGroups” tool from Picard (v 2.7.0-SNAPSHOT; http://broadinstitute.github.io/picard/). We then ran all BAM files through GATK’s “ASEReadCounter” v 3.6-0-g89b7209 (17) with default options to obtain read counts for each allele. This step was carried out once on all 6 North American samples, once on all 4 SB/Sb Taiwanese samples and once on all 24 South American samples, using the fixed differences between variants generated previously for each population. In both cases we used the original genome of *S. invicta* (version gnG; RefSeq GCF_000188075.1) as a reference.

We then imported the resulting allele-specific SNP read counts per sample generated by GATK into R v3.4.4 (46). We used the R packages “GenomicRanges” v1.26.4 (82) and “GenomicFeatures” v1.26.3 (82) along with the NCBI protein-coding gene annotation for *S. invicta* (gnG assembly, release 100) to identify which SNPs are in which genes. This is because allele-specific expression of long genes with several fixed SNPs between variants could be overestimated if the reads per SNP are counted individually. To avoid this, we estimated the total expression level for a particular allele (*i.e.*, the SB or Sb variant for any given gene) as the median of all SNP-specific read counts per gene and per variant. For instance, consider a gene with 3 fixed SNPs between SB and Sb for which the SB variants have support from 12, 15 and 18 reads, and the Sb variants from 5, 8 and 6 reads. In this particular case, we would report that the SB variant for this particular gene has an expression level of 15 reads and the Sb variant, 6 reads. If instead of this approach, we randomly select one of the possible SNPs for every gene, we find qualitatively similar results to those reported.Additionally, to test whether we would be able to detect allele-specific expression changes across body parts and castes in the South American data, we calculated allele-specific expression in the whole genome as a positive control. We used the VCF file containing all SNPs in the 26 males collected from South America. We retained only SNPs with expression data in all samples and a median of at least 1-x RNA coverage in each allele across all samples. After filtering, 1096 SNPs remained on which we were able to test for allele-specific expression. We then performed an allele specific expression analysis throughout the whole genome using body part and caste information from South American populations. Unlike the analysis of genes in the supergene region, in the whole genome analysis we cannot ensure that every individual sequenced is heterozygous for all SNPs. Indeed, the average frequency in the population for all the alleles analyzed was 0.41 with a standard deviation of ± 0.2. This implies that both alleles were not necessarily present in all samples. We therefore had far less power to detect allele-specific expression across body parts using data from the whole genome than using SNPs from the supergene region only. Despite this lack of power, we were able to detect significant (Wald test BH adjusted p<0.05) allele-specific expression changes across body parts of queens and workers in 15 SNPs. These significant SNPs were distributed across 9 genomic scaffolds. The significant differences in allele-specific expression were between a queen body part and whole bodies of workers.

### Identifying expression differences between the SB and Sb variants

We imported the estimated read counts generated by Kallisto into R using Tximport v1.2.0 (83) and DESeq2 v1.14.1 (43). For every sample, read counts for the SB alleles and for the Sb alleles come from a single sequencing library, thus standard normalization methods (84) are not applicable. As recommended by the developers of DESeq2 (85), we thus deactivated normalization by setting SizeFactors=1. For the North American and Taiwanese datasets (41), we only considered genes expressed in all samples for downstream analyses, whereas for the South American populations RNA dataset, we only analyzed genes expressed in all replicates of at least one body part.

To have the strongest possible analysis of expression between the SB and Sb variants of the supergene region, we performed a joint analysis of RNAseq data from Taiwanese, South and North American populations. The South American dataset includes body part information, which is absent in the North American dataset. If both datasets were analyzed together with any the standard tools for gene expression analysis, the effect in expression levels arising from different body parts would be confounded with that arising from differences between the two datasets. To overcome this issue, we applied a linear mixed effects model on the log2 of the expression ratios between SB and Sb across populations and body parts, using the R packages lme4 v1.1-18.1 (86) and lmerTest v3.1-0 (87). The aim of this model was to identify the expression differences between SB and Sb across the different datasets, accounting for the proportion of variance explained by body part and population. We fitted the log2 expression ratios using a 0 intercept with gene, population and their interaction as fixed effects, and the interaction between gene and body part as random effects (formula: log2 expression ratio ∼ 0 + gene * population + (1|body part:gene)).

We also performed an additional linear mixed-effects model to test the effect of geography and ancestry on the allele-specific expression patterns within the supergene. We grouped the populations by geographical proximity (North and South America vs Taiwan) or by phylogenetic proximity (Taiwan and North America vs South America). We then fitted the log2 expression ratios using a 0 intercept, the main effects of gene, ancestry and geographic proximity and the interactions between ancestry and gene and between geographic proximity and gene as fixed effects. As random effects we used again the interaction between gene and body part (formula: log2 expression ratio ∼ 0 + gene * ancestry + gene * geography + (1| body_part:gene)). We then performed an analysis of variance on the model to estimate the size effects of each term.

For both models, the log2 expression ratios were weighed by a function of the total read counts per gene to reduce the impacts of genes with low expression which have extremely high variance. Here we only report the results of the fixed effects per gene after adjustment of p values for multiple testing following the Benjamini-Hochberg approach (88). For this joint analysis we only used genes that had fixed differences between SB and Sb in all three populations.

We additionally analyzed the allele-specific expression patterns between the SB and Sb variants of the supergene in each population independently using DESeq2 (43) as suggested by Castel and collaborators (79). The model formula used for the South American RNA-seq data used “body part” and “colony of origin” as blocking factors, and allele-specific expression, *i.e.* “variant effect”, as variable of interest. This analysis allowed us to detect differences in expression between variants specific to body part. Preliminary analyses showed that the interaction between “variant effect” and “body part” had no significant effect in any of the genes, and consequently, only the main “variant effect” was considered as the factor of interest for this analysis. The model formula for both the Wurm *et al.* (41) and the Fontana *et al.* (42) RNA-seq datasets included only whole bodies of queens. We thus used “sample” as a blocking factor and “variant effect” as variable of interest.

In all analyses, we report gene differences between variants as log2 expression ratios between the SB and the Sb counts. That is, genes with expression biased towards SB will produce positive log2 expression ratios whereas those biased towards Sb will produce a negative value. To check whether there was an overall bias towards either variant, we tested the significance of the deviation from 0 for the median log2 expression ratios between SB and Sb via a Wilcoxon sum rank test. Significant differences between alleles for genes common in both populations are reported in Figure 1, we report the results for all genes of each independent analysis in Figure1-Figure supplement 2 and Table S3a for South American populations, in Figure1-Figure supplement 3 and Table S3b for North American populations and in Figure1-Figure supplement 4 and Table S3c for Taiwanese populations.

### Expression differences between social forms

We determined the expression levels for all samples from the North American populations (45) by using the count mode in Kallisto v0.44.0 (89) using S. invicta coding sequences (version gnG; RefSeq GCF_000188075.1, release 100) as a reference. We imported the estimated counts into DESeq2 v1.14.1 (43) using Tximport v1.2.0 (83). We compared the DESeq2 normalized expression levels between social forms, determining significance of differential expression using the default Wald test for pairwise comparisons between genes. For each caste we estimated the proportion of significantly differentially to non-differentially expressed genes within and outside the supergene region based on supergene region coordinates from Pracana et al. (34). We then used the R packages GenomicRanges and GenomicFeatures (82) along with the annotations of *S. invicta* coding sequences to locate each gene with expression information in the genome. Our analyses are restricted to the 10,481 known *S. invicta* genes that can be reliably placed within or outside the supergene region; other genes are on scaffolds which lack chromosomal locations.

### Expression differences between variants and social forms

We fitted a model to test whether there is a significant relationship between allele-specific expression differences between supergene variants in the Wurm *et al* dataset (41), and gene expression differences between social forms (log2 expression ratios using the Morandin *et al* dataset (45)). We examined the overall trend in allele-specific expression patterns within the supergene (*i.e.*, any bias towards expression of either the SB or Sb allelic variant). We predicted that there would be significant association between allele-specific expression and the expression differences between social forms. In particular, assuming evolutionary conflict between the social forms, we predicted that genes which are relatively highly expressed in multiple-queen colonies would have relatively high expression of the Sb allele, whereas those genes with relatively low expression in multiple-queen colonies would have relatively high expression of the SB allele.

We obtained relative expression levels using DESeq2 for both comparisons: single-queen *vs.* multiple-queen expression for each gene (*X*_*SQ*_ *vs X*_*MQ*_) from the Wurm *et al* dataset (41) and expression of the SB allelic variant *vs*. the Sb (*X*_*B*_ *vs. X*_*b*_) within each gene from the Morandin *et al* dataset (45) DESeq2 returned an estimate of log2(X_B_ / X_b_) for the allele specific expression and log2(X_SQ_ / X_MQ_) for the differences among colony types. We transformed these to relative proportions of gene expression *P*_*B*_ = *X*_*B*_ / (*X*_*B +*_ *X*_*b*_) and *P*_*MQ*_ = *X*_*MQ*_ / (*X*_*SQ*_ + *X*_*MQ*_).

## Acknowledgments

This research was possible thanks to the funding provided by the Natural Environment Research Council (NE/L00626X/1 and NERC EOS Cloud to YW; NE/L002485/1 to CM-R), Deutscher Akademischer Austauschdienst (DAAD) Postdoc Program (570704 83 to E.S.); European Commission Marie Curie Actions (PIEF-GA-2013-623713 to ES and YW); Biotechnology and Biological Sciences Research Council (BB/ K004204/1 to YW). We thank Emiliano Boné for help with ant rearing, Claudia Castillo-Carrillo for help during sample collections, Dr. Monika Struebig for preparing RNA-seq libraries, Phillip Howard, Dr. Chloe Economou and Martin Tran for their technical wet lab support, Wurm lab members and colleagues in the Department of Organismal Biology for valuable input and discussions, and the ITS Research Group and Adrian Lärkeryd at Queen Mary University of London for computational support and access to the Apocrita High Performance Computing facility (http://doi.org/10.5281/zenodo.438045) and NERC EOS Cloud.

## Data availability

We deposited the genomic and transcriptomic reads we generated from South American *Solenopsis invicta* on NCBI SRA (PRJNA542606). All analysis scripts used will be made available at http://github.com/ MartinezRuiz-Carlos/2019-11_allele_specific_expression_fire_ant upon publication. During review they are available at http://bit.ly/ase_code.

## Author Contributions

YW, RAN and CM-R conceived and designed the study; CM-R extracted the RNA samples and performed all the gene and allele expression analyses; RP analyzed the DNA data and performed the variant calling and filtering; ES and CIP collected the samples in the field; ES extracted DNA; CM-R, RAN and YW drafted the manuscript and all authors contributed to later versions of the manuscript.

## Figure supplements

**Figure 1-Figure supplement 1.**
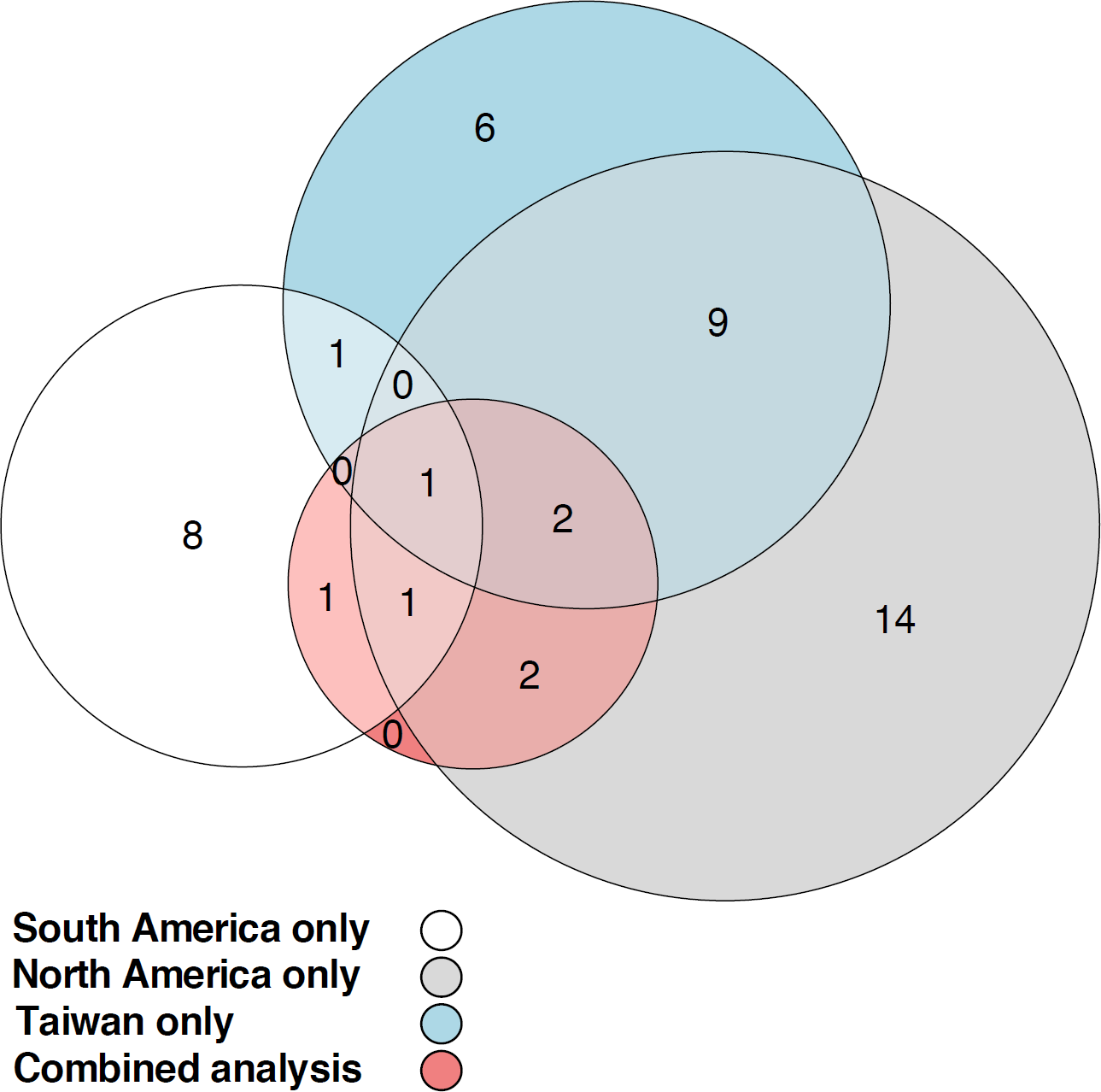
Euler diagram showing the overlapping number of genes with allele-specific expression according to four different comparisons: using combined data from Taiwanese, North and South American populations (red circle), using data from South American populations only (white circle), from North American populations only (grey circle) or from Taiwanese populations only (light blue circle). The combined analysis detected 7 genes with allele-specific expression across both populations, 3 of which were independently detected using only South American populations, 6 using only North American populations and 3 using only Taiwanese populations.

**Figure 1-Figure supplement 2.**
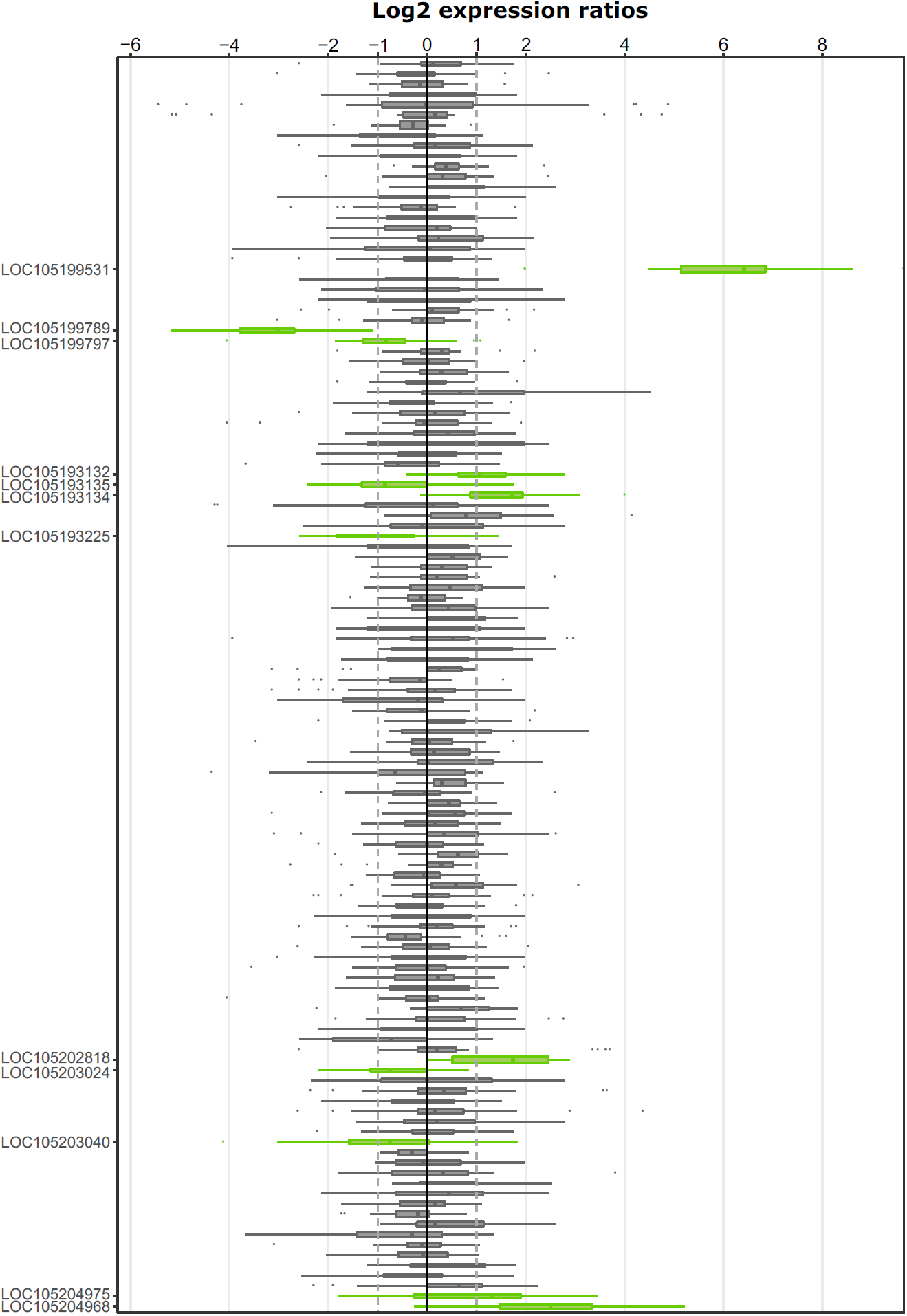
Allele-specific expression for genes in the fire ant social supergene. Differences in allele-specific expression between variants (x axis) for all genes in the supergene with enough expression information (y axis) for South American samples (information from body parts of queens and whole bodies of workers merged together). Significant expression differences (BH adjusted p<0.05 from Wald test in DESeq2) are indicated by green boxes and the RefSeq ID of the significant gene is outlined in the y axis. Non-significant differences are marked by grey boxes. Within each plot, each box shows the distribution of log2 expression ratios between SB and Sb. Genes with values above the solid line (log2 expression ratios=0) are more highly expressed in SB and below, they are more highly expressed in Sb. The dashed line shows log2 expression ratios=1, i.e., a two-fold gene expression difference in either direction. Genes are in chromosomal order. The relative size of each box within either plot is correlated with the median read count in which the log2 expression ratios are based (the thicker the box, the more read counts that particular gene has).

**Figure 1-Figure supplement 3.**
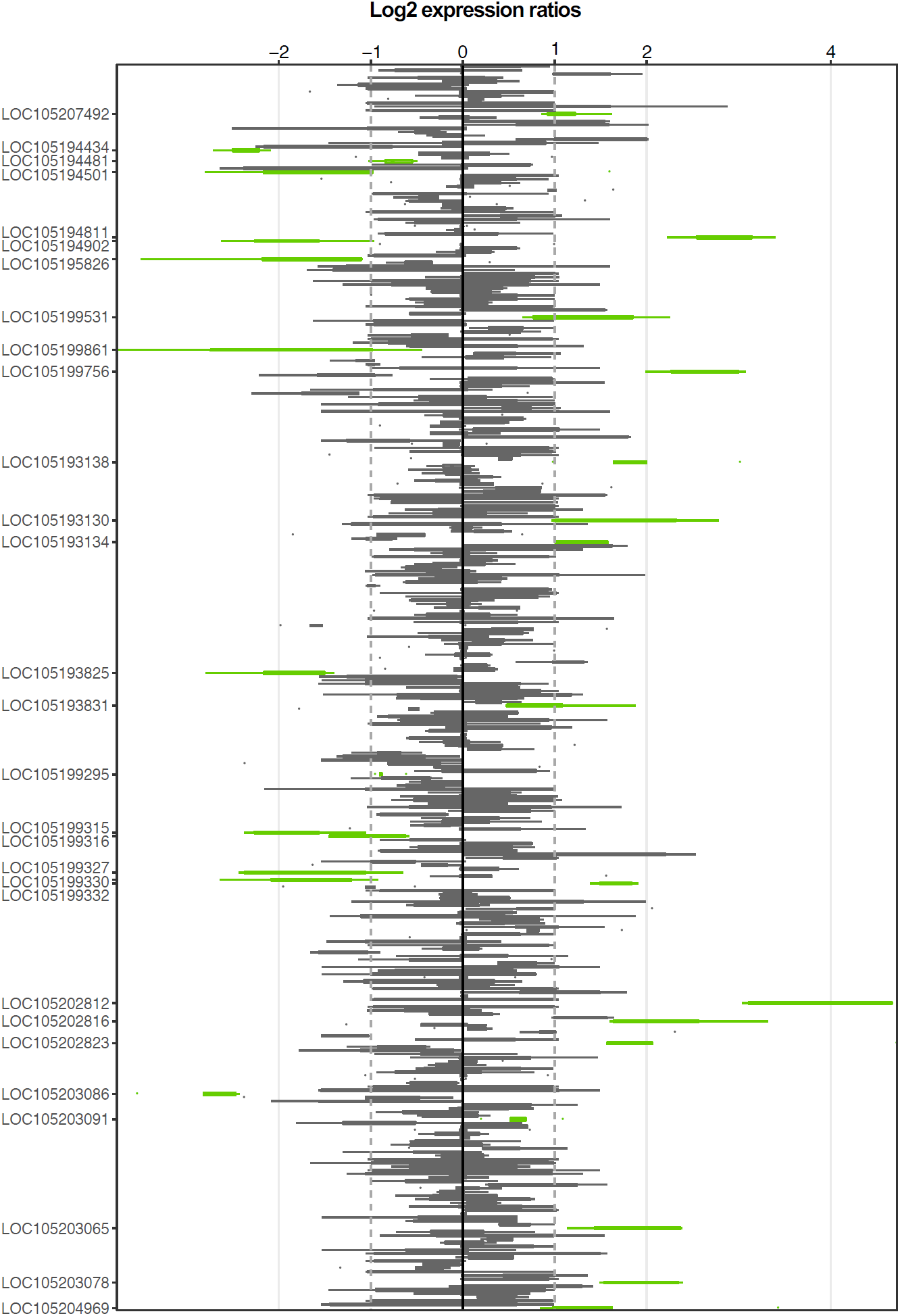
Allele-specific expression for genes in the fire ant social supergene. Differences in allele-specific expression between variants (x axis) for all genes in the supergene with enough expression information (y axis) for North American queens (whole body). Significant expression differences (BH adjusted p<0.05 from Wald test in DESeq2) are indicated by green boxes and the RefSeq ID of the significant gene is outlined in the y axis. Non-significant differences are marked by grey boxes. Within each plot, each box shows the distribution of log2 expression ratios between SB and Sb. Genes with values above the solid line (log2 expression ratios=0) are more highly expressed in SB and below, they are more highly expressed in Sb. The dashed line shows log2 expression ratios=1, i.e., a two-fold gene expression difference in either direction. Genes are in chromosomal order. The relative size of each box within either plot is correlated with the median read count in which the log2 expression ratios are based (the thicker the box, the more read counts that particular gene has).

**Figure 1-Figure supplement 4.**
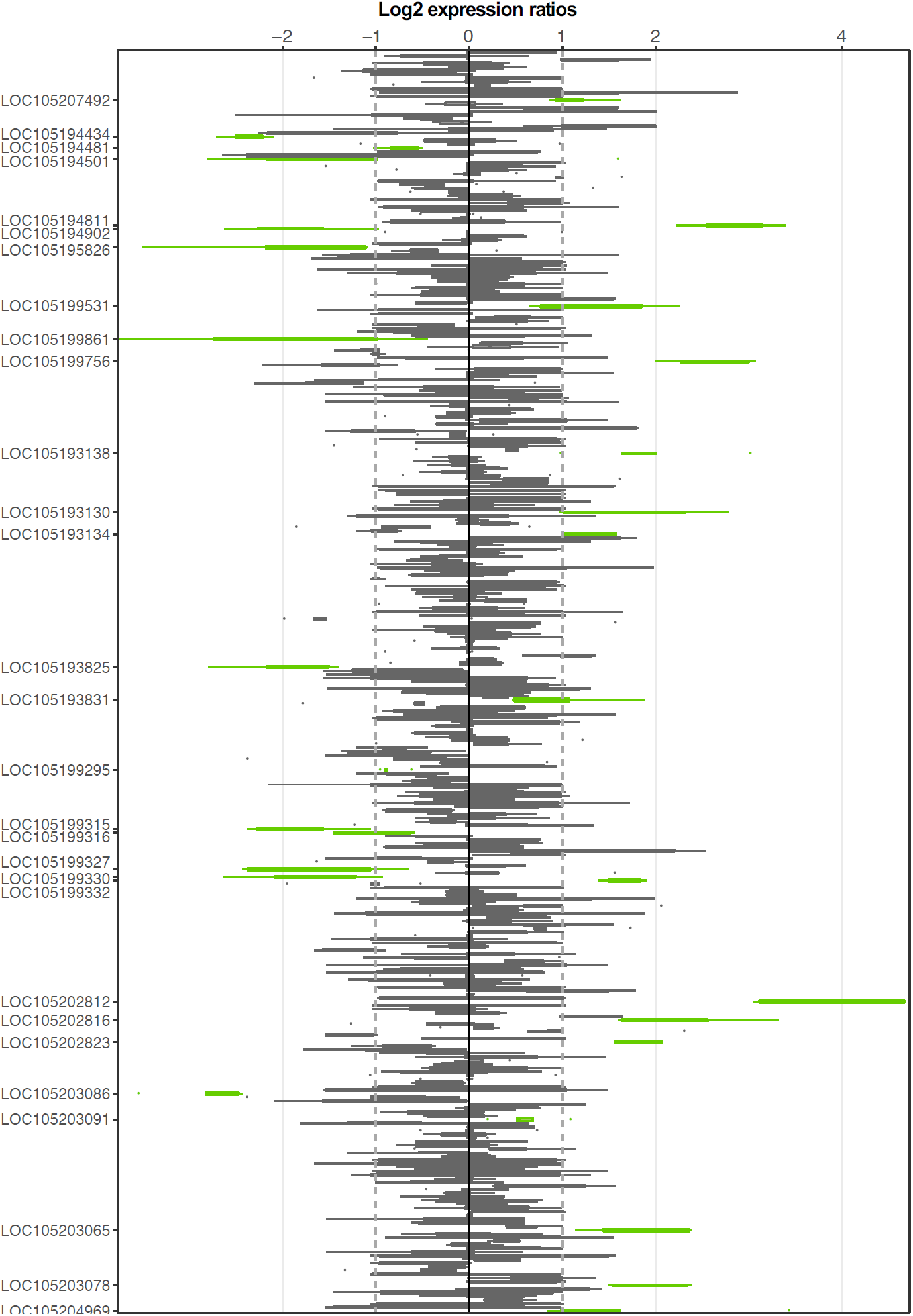
Allele-specific expression for genes in the fire ant social supergene. Differences in allele-specific expression between variants (x axis) for all genes in the supergene with enough expression information (y axis) for Taiwanese queens (whole body). Significant expression differences (BH adjusted p<0.05 from Wald test in DESeq2) are indicated by green boxes and the RefSeq ID of the significant gene is outlined in the y axis. Non-significant differences are marked by grey boxes. Within each plot, each box shows the distribution of log2 expression ratios between SB and Sb. Genes with values above the solid line (log2 expression ratios=0) are more highly expressed in SB and below, they are more highly expressed in Sb. The dashed line shows log2 expression ratios=1, i.e., a two-fold gene expression difference in either direction. Genes are in chromosomal order. The relative size of each box within either plot is correlated with the median read count in which the log2 expression ratios are based (the thicker the box, the more read counts that particular gene has).

**Figure 1-Figure supplement 5.**
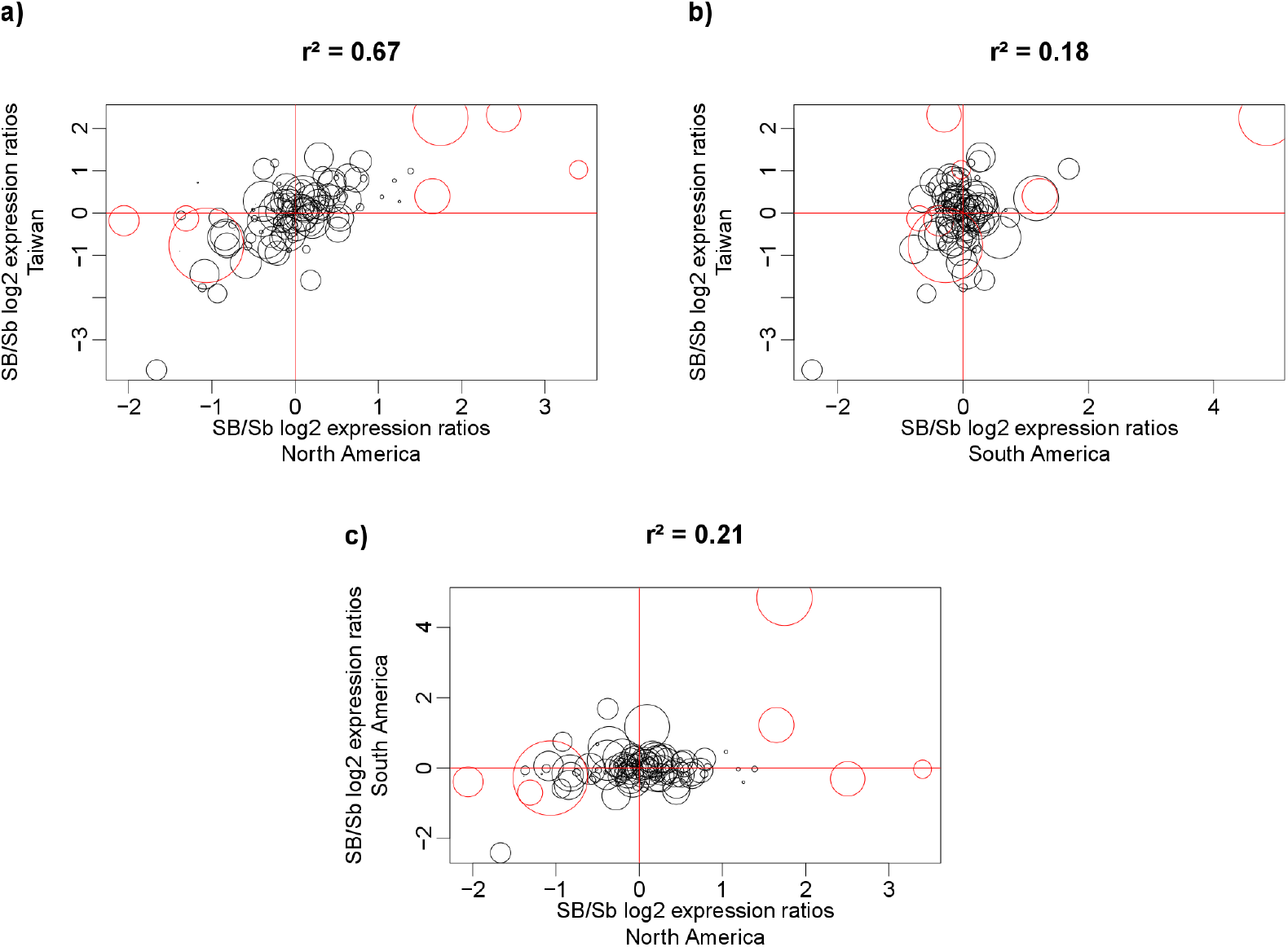
Correlation of log2 allele-specific expression ratios between the SB and Sb variants in heterozygous queens from three populations: South American data we generated, North American data (from Morandin *et al.* 2016), and Taiwanese data (from Fontana *et al.* 2020). We show correlations: **a)** between Taiwanese and North American populations; **b)** between Taiwanese and South American populations; and **c)** between South American and North American populations. Positive values represent higher expression of the SB allele, negative values represent higher expression of the Sb allele. Each dot represents a single gene within the supergene region. The size of the dots are proportional to the average expression level of that gene. Red dots represent the genes detected by the linear mixed effects model as significantly differentially expressed between SB and Sb across populations. The correlation r^2^ between each pair of populations was calculated using the Spearman method, with each gene being weighted by mean expression level (read counts).

**Figure 3-Figure supplement 1.**
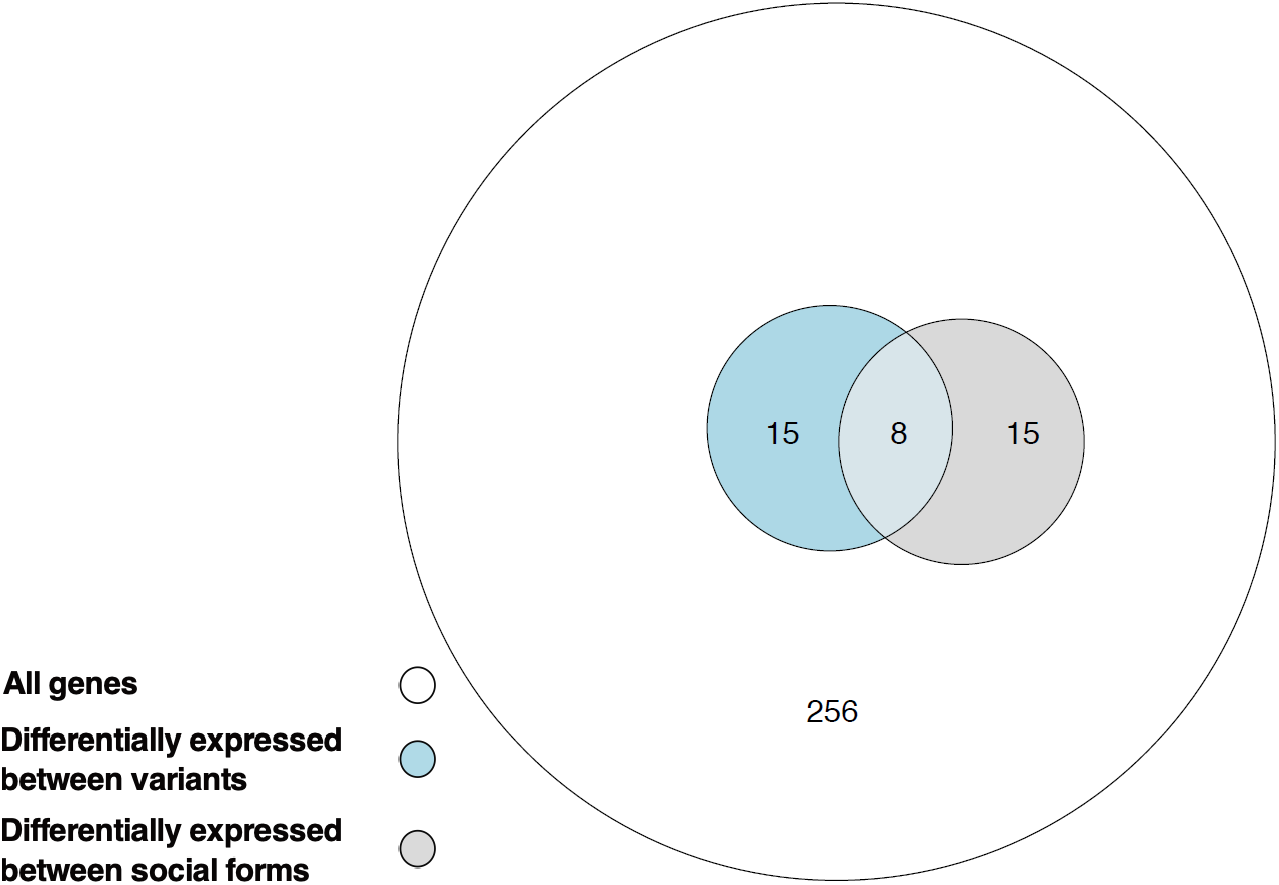
Euler diagram showing the overlap of genes with expression differences between variants (light blue circle) and social forms (grey circle) out of all genes within the supergene region with expression information in both comparisons on the North American dataset (white circle).

## References

1. D. Charlesworth, The status of supergenes in the 21st century: recombination suppression in Batesian mimicry and sex chromosomes and other complex adaptations. Evol. Appl. 9, 74–90 (2016).

2. C. D. Darlington, K. Mather, The elements of genetics (George Allen & Unwin Ltd: London, 1949).

3. M. J. Thompson, C. D. Jiggins, Supergenes and their role in evolution. Heredity 113, 1–8 (2014).

4. J. Li, et al., Genetic architecture and evolution of the S locus supergene in *Primula vulgaris*. Nat. Plants. 2 (2016).

5. S. Branco, et al., Multiple convergent supergene evolution events in mating-type chromosomes. Nat. Commun. 9, 2000 (2018).

6. M. Joron, et al., Chromosomal rearrangements maintain a polymorphic supergene controlling butterfly mimicry. Nature 477, 203–206 (2011).

7. K. Kunte, et al., doublesex is a mimicry supergene. Nature 507, 229–232 (2014).

8. W. M. Zinzow-Kramer, et al., Genes located in a chromosomal inversion are correlated with territorial song in white-throated sparrows. Genes Brain Behav. 14, 641–654 (2015).

9. E. M. Tuttle, et al., Divergence and functional degradation of a sex chromosome-like supergene. Curr. Biol. 26, 344–350 (2016).

10. C. Küpper, et al., A supergene determines highly divergent male reproductive morphs in the ruff. Nat. Genet. 48, 79–83 (2016).

11. S. Lamichhaney, et al., Structural genomic changes underlie alternative reproductive strategies in the ruff (*Philomachus pugnax*). Nat. Genet. 48, 84–88 (2016).

12. R. Bergero, D. Charlesworth, The evolution of restricted recombination in sex chromosomes. Trends Ecol. Evol. 24, 94–102 (2009).

13. A. E. Wright, et al., Convergent recombination suppression suggests role of sexual selection in guppy sex chromosome formation. Nat. Commun. 8, 14251 (2017).

14. J. E. Mank, The transcriptional architecture of phenotypic dimorphism. Nat. Ecol. Evol. 1, 6 (2017).

15. N. Zemp, et al., Evolution of sex-biased gene expression in a dioecious plant. Nat. Plants 2, 16168 (2016).

16. B. Vicoso, V. B. Kaiser, D. Bachtrog, Sex-biased gene expression at homomorphic sex chromosomes in emus and its implication for sex chromosome evolution. Proc. Natl. Acad. Sci. U. S. A. 110, 6453–6458 (2013).

17. P. P. Khil, N. A. Smirnova, P. J. Romanienko, R. D. Camerini-Otero, The mouse X chromosome is enriched for sex-biased genes not subject to selection by meiotic sex chromosome inactivation. Nat. Genet. 36, 642–646 (2004).

18. J. Parsch, H. Ellegren, The evolutionary causes and consequences of sex-biased gene expression. Nat. Rev. Genet. 14, 83–87 (2013).

19. C. Dufresnes, et al., Sex-chromosome homomorphy in palearctic tree frogs results from both turnovers and X–Y recombination. Mol. Biol. Evol. 32, 2328–2337 (2015).

20. M. Nozawa, N. Fukuda, K. Ikeo, T. Gojobori, Tissue- and stage-dependent dosage compensation on the neo-X chromosome in *Drosophila pseudoobscura*. Mol. Biol. Evol. 31, 614–624 (2014).

21. A. A. Alekseyenko, et al., Conservation and *de novo* acquisition of dosage compensation on newly evolved sex chromosomes in *Drosophila*. Genes Dev. 27, 853–858 (2013).

22. A. Muyle, et al., Rapid de novo evolution of X chromosome dosage compensation in *Silene latifolia*, a plant with young sex chromosomes. PLoS Biol. 10, e1001308 (2012).

23. D. Charlesworth, B. Charlesworth, G. Marais, Steps in the evolution of heteromorphic sex chromosomes. Heredity 95, 118–128 (2005).

24. M. Stöck, et al., Ever-young sex chromosomes in European tree frogs. PLoS Biol. 9, e1001062 (2011).

25. E. Cavoto, S. Neuenschwander, J. Goudet, N. Perrin, Sex-antagonistic genes, XY recombination and feminized Y chromosomes. J. Evol. Biol. 31, 416–427 (2018).

26. S. Branco, et al., Evolutionary strata on young mating-type chromosomes despite the lack of sexual antagonism. Proc. Natl. Acad. Sci. U. S. A. 114, 7067–7072 (2017).

27. L. Gu, J. R. Walters, Evolution of sex chromosome dosage compensation in animals: A beautiful theory, undermined by facts and bedeviled by details. Genome Biol. Evol. 9, 2461–2476 (2017).

28. C. Ellison, D. Bachtrog, Recurrent gene co-amplification on *Drosophila* X and Y chromosomes. PLoS Genet. 15, e1008251 (2019).

29. M. Cowley, R. J. Oakey, Transposable elements re-wire and fine-tune the transcriptome. PLoS Genet. 9, e1003234 (2013).

30. D. Bachtrog, Y-chromosome evolution: emerging insights into processes of Y-chromosome degeneration. Nat. Rev. Genet. 14, 113–124 (2013).

31. J. Wang, et al., A Y-like social chromosome causes alternative colony organization in fire ants. Nature 493, 664–668 (2013).

32. D. Gotzek, K. G. Ross, Genetic regulation of colony social organization in fire ants: an integrative overview. Q. Rev. Biol. 82, 201–226 (2007).

33. G. N. Fritz, R. K. Vander Meer, C. A. Preston, Selective male mortality in the red imported fire ant, *Solenopsis invicta*. Genetics 173, 207–213 (2006).

34. R. Pracana, A. Priyam, I. Levantis, R. A. Nichols, Y. Wurm, The fire ant social chromosome supergene variant Sb shows low diversity but high divergence from SB. Mol. Ecol. 26, 2864–2879 (2017).

35. E. Stolle, et al., Degenerative expansion of a young supergene. Mol. Biol. Evol. 36, 553–561 (2019).

36. Y.-C. Huang, V. D. Dang, N.-C. Chang, J. Wang, Multiple large inversions and breakpoint rewiring of gene expression in the evolution of the fire ant social supergene. Proc. Biol. Sci. 285 (2018).

37. I. Darolti, et al., Extreme heterogeneity in sex chromosome differentiation and dosage compensation in livebearers. Proc. Natl. Acad. Sci. U. S. A. 116, 19031–19036 (2019).

38. R. Pracana, et al., Fire ant social chromosomes: Differences in number, sequence and expression of odorant binding proteins. Evol. Lett. 1, 199–210 (2017).

39. D. W. Hall, M. A. D. Goodisman, The effects of kin selection on rates of molecular evolution in social insects. Evolution 66, 2080–2093 (2012).

40. K. G. Ross, D. D. Shoemaker, Estimation of the number of founders of an invasive pest insect population: the fire ant *Solenopsis invicta* in the USA. Proc. Biol. Sci. 275, 2231–2240 (2008).

41. Y. Wurm, et al., The genome of the fire ant *Solenopsis invicta*. Proc. Natl. Acad. Sci. U. S. A. 108, 5679–5684 (2011).

42. S. Fontana, et al., The fire ant social supergene is characterized by extensive gene and transposable element copy number variation. Mol. Ecol. 29, 105–120 (2020).

43. M. I. Love, H. Wolfgang, A. Simon, Moderated estimation of fold change and dispersion for RNA-seq data with DESeq2. Genome Biol. 15 (2014).

44. M. S. Ascunce, et al., Global invasion history of the fire ant *Solenopsis invicta*. Science 331, 1066–1068 (2011).

45. C. Morandin, et al., Comparative transcriptomics reveals the conserved building blocks involved in parallel evolution of diverse phenotypic traits in ants. Genome Biol. 17, 43 (2016).

46. R Core Team, R: A language and environment for statistical computing. R Foundation for Statistical Computing, Vienna, Austria (2017).

47. J. E. Mank, N. Wedell, D. J. Hosken, Polyandry and sex-specific gene expression. Philos. Trans. R. Soc. Lond. B Biol. Sci. 368, 20120047 (2013).

48. Y.-C. Huang, J. Wang, Did the fire ant supergene evolve selfishly or socially? Bioessays 36, 200–208 (2014).

49. D. R. Denver, et al., The transcriptional consequences of mutation and natural selection in *Caenorhabditis elegans*. Nat. Genet. 37, 544–548 (2005).

50. S. A. Rifkin, D. Houle, J. Kim, K. P. White, A mutation accumulation assay reveals a broad capacity for rapid evolution of gene expression. Nature 438, 220–223 (2005).

51. E. Stolle, R. Pracana, Y. Wurm, Degenerative expansion of a young supergene. Mol. Biol. Evol. 36, 553–561 (2019).

52. M. M. Patten, The X chromosome favors males under sexually antagonistic selection. Evolution 73, 84–91 (2019).

53. D. Sun, I. Huh, W. M. Zinzow-Kramer, D. L. Maney, S. V. Yi, Rapid regulatory evolution of a nonrecombining autosome linked to divergent behavioral phenotypes. Proc. Natl. Acad. Sci. U. S. A. 115, 2794–2799 (2018).

54. J. E. Mank, Sex chromosome dosage compensation: definitely not for everyone. Trends Genet. 29, 677–683 (2013).

55. L. Keller, K. G. Ross, Selfish genes: a green beard in the red fire ant. Nature 394, 573 (1998).

56. M. J. B. Krieger, K. G. Ross, Identification of a major gene regulating complex social behavior. Science 295, 328–332 (2002).

57. M. Laturney, J.-C. Billeter, Neurogenetics of female reproductive behaviors in *Drosophila melanogaster*. Adv. Genet. 85, 1–108 (2014).

58. J. Bohbot, F. Sobrio, P. Lucas, P. Nagnan-Le Meillour, Functional characterization of a new class of odorant-binding proteins in the moth *Mamestra brassicae*. Biochem. Biophys. Res. Commun. 253, 489–494 (1998).

59. F. Wolschin, H. Shpigler, G. V. Amdam, G. Bloch, Size-related variation in protein abundance in the brain and abdominal tissue of bumble bee workers. Insect Mol. Biol. 21, 319–325 (2012).

60. K. A. Yoon, et al., Comparative functional venomics of social hornets *Vespa crabro* and *Vespa analis*. J. Asia. Pac. Entomol. 18, 815–823 (2015).

61. M. M. Steller, S. Kambhampati, D. Caragea, Comparative analysis of expressed sequence tags from three castes and two life stages of the termite *Reticulitermes flavipes*. BMC Genomics 11, 463 (2010).

62. P. Pelosi, I. Iovinella, J. Zhu, G. Wang, F. R. Dani, Beyond chemoreception: diverse tasks of soluble olfactory proteins in insects. Biol. Rev. Camb. Philos. Soc. 93, 184–200 (2018).

63. J. T. Robinson, et al., Integrative genomics viewer. Nat. Biotechnol. 29, 24–26 (2011).

64. B. R. Johnson, J. Atallah, D. C. Plachetzki, The importance of tissue specificity for RNA-seq: highlighting the errors of composite structure extractions. BMC Genomics 14, 586 (2013).

65. M. Martin, Cutadapt removes adapter sequences from high-throughput sequencing reads. EMBnet.journal 17, 10–12 (2011).

66. A. Dobin, et al., STAR: ultrafast universal RNA-seq aligner. Bioinformatics 29, 15–21 (2013).

67. O. Tange, GNU parallel-the command-line power tool. The USENIX Magazine 36, 42–47 (2011).

68. T. King, S. Butcher, L. Zalewski, Apocrita—high performance computing cluster for Queen Mary University of London. Zenodo (2017).

69. P. Ewels, M. Magnusson, S. Lundin, M. Käller, MultiQC: summarize analysis results for multiple tools and samples in a single report. Bioinformatics 32, 3047–3048 (2016).

70. L. Wang, et al., Measure transcript integrity using RNA-seq data. BMC Bioinformatics 17, 58 (2016).

71. K. G. Ross, M. J. B. Krieger, L. Keller, D. D. Shoemaker, Genetic variation and structure in native populations of the fire ant *Solenopsis invicta*: evolutionary and demographic implications. Biol. J. Linn. Soc. Lond. 92, 541–560 (2007).

72. M. E. Ahrens, K. G. Ross, D. D. Shoemaker, Phylogeographic structure of the fire ant *Solenopsis invicta* in its native South American range: Roles of natural barriers and habitat connectivity. Evolution 59, 1733–1743 (2005).

73. B. Langmead, C. Trapnell, M. Pop, S. L. Salzberg, Ultrafast and memory-efficient alignment of short DNA sequences to the human genome. Genome Biol. 10, R25 (2009).

74. E. Garrison, G. Marth, Haplotype-based variant detection from short-read sequencing. ArXiv preprint 1207.3907 (2012).

75. H. Li, et al., The Sequence Alignment/Map format and SAMtools. Bioinformatics 25, 2078–2079 (2009).

76. V. Obenchain, et al., VariantAnnotation: a Bioconductor package for exploration and annotation of genetic variants. Bioinformatics 30, 2076–2078 (2014).

77. P. Danecek, et al., The variant call format and VCFtools. Bioinformatics 27, 2156–2158 (2011).

78. P. Cingolani, et al., A program for annotating and predicting the effects of single nucleotide polymorphisms, SnpEff: SNPs in the genome of *Drosophila melanogaster* strain w1118; iso-2; iso-3. Fly 6, 80–92 (2012).

79. S. E. Castel, A. Levy-Moonshine, P. Mohammadi, E. Banks, T. Lappalainen, Tools and best practices for data processing in allelic expression analysis. Genome Biol. 16, 195 (2015).

80. B. van de Geijn, G. McVicker, Y. Gilad, J. K. Pritchard, WASP: allele-specific software for robust molecular quantitative trait locus discovery. Nat. Methods 12, 1061–1063 (2015).

81. A. McKenna, et al., The genome analysis toolkit: A MapReduce framework for analyzing next-generation DNA sequencing data. Genome Res. 20, 1297–1303 (2010).

82. M. Lawrence, et al., Software for computing and annotating genomic ranges. PLoS Comput. Biol. 9, e1003118 (2013).

83. C. Soneson, M. I. Love, M. D. Robinson, Differential analyses for RNA-seq: transcript-level estimates improve gene-level inferences. F1000Res. 4, 1521 (2015).

84. M.-A. Dillies, et al., A comprehensive evaluation of normalization methods for Illumina high-throughput RNA sequencing data analysis. Brief. Bioinform. 14, 671–683 (2013).

85. M. I. Love, Using RNA-seq DE methods to detect allele-specific expression (2017) URL:http://rpubs.com/mikelove/ase (accessed May 22, 2018).

86. D. Bates, M. Mächler, B. Bolker, S. Walker, Fitting Linear Mixed-Effects Models using lme4. ArXiv preprint 1406.5823 (2014).

87. A. Kuznetsova, P. B. Brockhoff, R. H. B. Christensen, lmerTest package: tests in linear mixed effects models. J. Stat. Softw. 82 (2017).

88. Y. Benjamini, Y. Hochberg, Controlling the False Discovery Rate: A Practical and Powerful Approach to Multiple Testing. J. R. Stat. Soc. Series B Stat. Methodol. 57, 289–300 (1995).

89. N. L. Bray, H. Pimentel, P. Melsted, L. Pachter, Near-optimal probabilistic RNA-seq quantification. Nat. Biotechnol. 34, 525–527 (2016).

